# Integrative single-cell characterization of frugivory adaptations in the bat kidney and pancreas

**DOI:** 10.1101/2023.02.12.528204

**Authors:** Wei E. Gordon, Seungbyn Baek, Hai P. Nguyen, Yien-Ming Kuo, Rachael Bradley, Alex Galazyuk, Insuk Lee, Melissa R. Ingala, Nancy B. Simmons, Tony Schountz, Lisa Noelle Cooper, Ilias Georgakopoulos-Soares, Martin Hemberg, Nadav Ahituv

## Abstract

Frugivory evolved multiple times in mammals, including bats. However, the cellular and molecular components driving it remain largely unknown. Here, we used integrative single-cell sequencing on insectivorous and frugivorous bat kidneys and pancreases and identified key cell population, gene expression and regulatory element differences associated with frugivorous adaptation that also relate to human disease, particularly diabetes. We found an increase in collecting duct cells and differentially active genes and regulatory elements involved in fluid and electrolyte balance in the frugivore kidney. In the frugivorous pancreas, we observed an increase in endocrine and a decrease in exocrine cells and differences in genes and regulatory elements involved in insulin regulation. Combined, our work provides novel insights into frugivorous adaptation that also could be leveraged for therapeutic purposes.

## Introduction

Frugivory, the ability to thrive on a mostly fruit diet, has allowed mammals to expand their ecological niches (*1*). In bats, frugivory has evolved independently from the insectivorous ancestor (*2*) in two separate families: Pteropodidae (Old World fruit bats; OWFBs) and Phyllostomidae (New World fruit bats; NWFBs) (*3*). Adaptation to a primarily fruit-rich diet requires multiple morphological and metabolic adaptations. In bats, these include sensory adaptations like enhanced vision (*4, 5*) and olfaction (*6, 7*), morphological adaptations including the cranium (*8–12*), tongue (*13*), small intestine (*14, 15*), kidney (*16, 17*), and pancreas (*18, 19*), and metabolic adaptations including transporter losses (*20, 21*) and increased transporter activity (*3, 16*).

One of the main challenges a fruit-centered diet poses is high blood sugar levels, which can lead to diabetes. To overcome this, an animal must rapidly control high sugar levels. Interestingly, although fruit bats consume more sugar than non-frugivorous bats, they can lower their blood sugar faster (*3, 22–24*). Fruit bats demonstrate high sensitivity to glucose and insulin (*19*) and can directly fuel their metabolic needs with exogenous sugars (*25*). The pancreas is responsible for generating hormones that regulate blood sugar and appetite, such as insulin and glucagon (*26*), as well as secreting enzymes and digestive juices into the small intestine (*27*). The high sensitivity to sugar levels in fruit bats is thought to be supported by an expansion of endocrine tissue in the pancreas (*18, 19*) and by loss of genes involved in insulin metabolism and signaling (*20*).

Fruit is rich in water and low in electrolytes like sodium and calcium. The kidney is responsible for maintaining water and salt balance, filtering the blood of waste, and maintaining blood pressure (*28*). The kidney is also involved in metabolizing sugars, generating glucose, and clearing insulin from circulation (*29*). Several kidney modifications evolved in fruit bats that lead to dilute urine production (*30*), including an increased renal cortex and a decreased renal medulla relative to insect-eating bats (*16, 17*) as well as several transporter losses (*20*).

How mammals adapted to new food sources at the molecular level remains largely unknown. Genetic investigations of insectivory have identified evolutionary adaptations of metabolic enzymes, including gene duplication events of chitin-hydrolyzing enzymes (*31, 32*). Comparative investigations of mammalian frugivorous adaptation have been primarily performed on a gene-by-gene basis, focusing on metabolism. These include genes such as *GYS1* and *GYS2* (glycogenesis), *NRF2* (antioxidant regulation), *TAT* (protein catabolism), *SLC2A4* (glucose transport), and *AGT* (glyoxylate detoxification) (*33–39*). In terms of gene regulatory elements, there are a limited number of studies. One example, is an 11 base pair (bp) deletion observed in both fruit bat families (OWFB and NWFB) in the proximal promoter of *SLC2A2*, which encodes the glucose transporter 2 (GLUT2), that was suggested to be responsible for the difference in liver *SLC2A2* expression in fruit bats (*3*). However, no systematic unbiased large-scale genomic studies have been performed to comprehensively identify the cellular and molecular factors governing frugivory adaptation.

Here, we used integrative single-cell RNA-seq and ATAC-seq on adult insectivorous (*Eptesicus fuscus*) and frugivorous (*Artibeus jamaicensis*) bats to identify cell populations, genes, and regulatory element differences that could be associated with frugivory adaptations in the kidney and pancreas. In total, we analyzed over 34,696 cells from eight insectivorous and seven frugivorous bats. We developed a cross-species integrated analysis framework for non-model organism genomes, and utilized human and mouse single-cell kidney and pancreas markers to annotate cell-types. Our data reveal cell-type, gene expression, and gene regulation differences between insectivorous and frugivorous bats. These include more collecting duct cells and gene expression changes involved in fluid and electrolyte balance in the frugivore kidney, and a relative decrease in exocrine cells and gene expression changes involved in insulin secretion and glucose response in the frugivore pancreas. Cell composition differences were further validated with immunofluorescence on insectivorous and frugivorous bat tissues, confirming a reduction in renal medulla and an expansion of endocrine tissue in fruit bats. Transcription factor (TF) analyses of single-cell ATAC-seq found divergent TF binding site (TFBS) usage between dietary phenotypes. Together, our single-cell multi-omics approach indicates that the frugivore kidneys and pancreases exhibit many signatures of diabetes, such as increased potassium secretion, gluconeogenesis, and glucose reabsorption in the kidney and hyperinsulinemia and hyperglycemia. In summary, our work provides the first single-cell datasets for bats that also compares kidneys and pancreases between closely related mammals of contrasting diets, providing insight into cell composition, gene expression, and gene regulation adaptations to frugivory that are also related to human disease, in particular with diabetes.

## Results

### Integrative single-cell profiling of insect- and fruit-eating bat kidneys and pancreases

To characterize the cell-types, genes and regulatory elements that differ between insectivorous and frugivorous bats, we conducted integrative single-cell sequencing (RNA-seq and ATAC-seq) on the kidneys and pancreases of four adult male insectivorous big brown bats (*Eptesicus fuscus*) (family: Vespertilionidae) and four adult male Jamaican fruit bats (*Artibeus jamaicensis*) (family: Phyllostomidae). The families of these bats diverged approximately 53.8 million years ago (Fig. 1A) (*40*). We used these bat species as they have publicly available genomes and are colonized in research labs. Big brown bats were fed a regular diet of mealworms in captivity, whereas Jamaican fruit bats were fed a variety of non-citrus fruits in captivity, such as cantaloupe and banana. We subjected these bats to an overnight fasting regime, followed by two big brown bats fed fruit-fed mealworms (to maximize fruit content) and two fruit bats fed fruit thirty minutes before euthanasia (Fig. 1B). We chose this time point because fruit bats digest food quickly and pass material within 30 minutes (*41*) and lower their blood sugar within 30 minutes (*3, 22–24*). Tissues were harvested immediately and flash-frozen. Nuclei were then isolated, subjected to fluorescence-activated cell sorting (FACS), and processed using the 10X Genomics Chromium single-cell Multiome ATAC + Gene Expression kit following established protocols (see Materials and Methods).

**Fig. 1.**
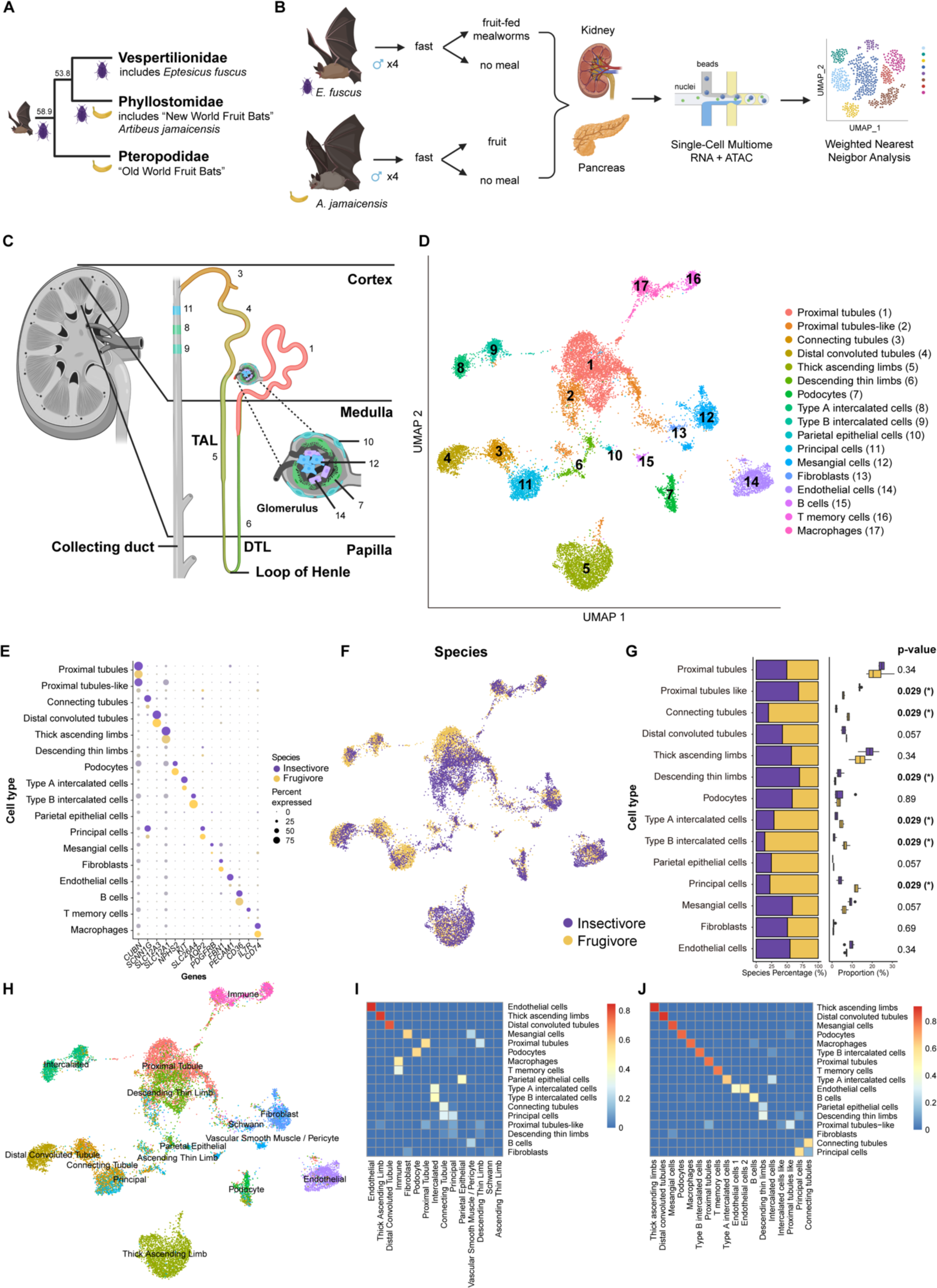
Joint scRNA and scATAC profiling of bat kidney. (**A**) Representative phylogenetic tree showing the evolution of frugivory in bats (order: Chiroptera). The beetle denotes lineage with insectivorous diet, and banana denotes lineage with frugivorous diet. Numbers are in millions of years (*40*). (**B**) Experimental design for joint scRNA and scATAC profiling of bat tissues (see Materials and Methods). (**C**) Diagram of kidney with zoom-in on the nephron. Colors correspond to cell-type colors in **D**. (**D**) Uniform Manifold Approximation and Projection (UMAP) of bat kidney cell-types based on scRNA-seq profiles. (**E**) Dot plot of marker gene expression across all bat kidney cell-types. Color intensity indicates the average expression level across all cells within a cell-type (purple or yellow is high; gray is low). (**F**) UMAP of bat kidney cell-types by species/dietary phenotype. (**G**) Plot of species percentage across all renal cells (left) with corresponding Wilcoxon rank-sum test for differential cell-type abundance (right) with significant p-values shown in bold font; * < 0.05. (**H**) UMAP of bat kidney cell-types annotated with human adult kidney single-cell reference data from Azimuth (*42*). (**I**) Jaccard overlap of annotations in **H** (horizontal) with our integrated species annotations in **D** (vertical). (**J**) Jaccard overlap of all annotations found in each species before integration (horizontal) with after integration in **D** (vertical).

As these bats are not widely used model organisms and have poorly assembled and annotated genomes, we made several modifications to analyze their multimodal data. These include (see Materials and Methods for more detail): 1) Removal of scaffolds < 50 kilo bases (kb) in length. Removal of these short scaffolds still allowed us to capture > 90% of the total sequences and genes for each genome (Fig. S1A). 2) Collapse of gene information in annotation files so that each gene is represented by a single “exon” transcript. 3) Due to technical reasons, ATAC-seq can contain a large number of mitochondrial reads (*43*), whose removal is needed for scATAC-seq analyses. While the Jamaican fruit bat has an annotated mitochondrial genome, the big brown bat does not. We thus used GetOrganelle (*44*) for assembly and MITOS (Bernt et al. 2013) for annotation to generate a big brown bat mitochondrial genome. Reassuringly, all mitochondrial genes identified in the Jamaican fruit bat genome were detected in the big brown bat genome. 4) To allow comparison of scRNA-seq and scATAC-seq between the two species, we used Orthofinder (*45*), using only one-to-one orthologues for subsequent comparisons to increase the number of shared features by 3.11% across the big brown bat genome and 3.24% across the Jamaican fruit bat genome.

We used cellranger-arc 2.0 (10X Genomics) to generate raw ATAC and RNA counts and Seurat (*42*) and Signac (*46*) for quality control and downstream analysis (Fig. S1B-C; Table S1). To merge samples, we created a common peak set across every sample within a species and within a tissue (Fig. S1D). To increase our sample sizes for phenotypic comparisons between species, we merged all samples within a species, hereinafter referred to as replicates. We combined two fasted and two fed insectivorous bat samples for each tissue (8,527 cells for kidney and 7,213 cells for pancreas), two fasted and two fed frugivorous bat samples for the kidney (9,315 cells), and two fasted and one fed fruit bat sample (due to one sample providing low quality sequencing data) for the pancreas (9,641 cells). To jointly analyze scRNA-seq and scATAC-seq within a species for each tissue, we used the R package Harmony (*47*) to correct for batch effects across replicates and applied “weighted nearest neighbor” analysis (*42*) (see Materials and Methods). To jointly analyze scRNA-seq and scATAC-seq across species for each tissue, we used gene activity scores to maximize the number of shared features for ATAC integration across species. We then employed the same methods used for analyzing within species as for cross-species analysis (Fig. S1D; see Materials and Methods). Cell-types were identified using canonical mouse and human markers. As bats diverged from human and mouse lineages approximately 75 million years ago (*48*), and our study presents the first functional genomics and gene expression dataset for bat kidney and pancreas, the number of shared markers per cell-type identified between bat, human and mouse was limited, restricting our ability to detect novel cell-types that could be unique to bats. Of note, while we observed some significant differences in gene expression between fasted and fed states in each species (Table S2-3), these were minimal compared to species gene expression differences (Fig. S2A-B) (longer treatments are likely needed for many transcriptional differences between fasted and fed (*49*)), with the exception of acinar cells, which are known to dominate total mRNA population from the pancreas (*50*) and permit rapid exocytosis of enzymes for digestion (*51*). Given our small sample sizes for treatment (N=1-2/treatment), we focused on subsequent analyses on species differences (N=3-4/species). We also did not observe significant differences in cell compositions between fasted and fed states in each species by co-varying neighborhood analysis (CNA) (Fig. S2C) (*52*), which was expected as cell-type differences are not likely to transpire following a 30 minute treatment.

### Insectivorous and frugivorous bat kidneys have different cell compositions

For the bat kidney, we initially annotated all major known cell-types using previously reported scRNA-seq markers from human and mouse kidney (*53–55*). These included proximal tubules (*CUBN*), connecting tubules (*SCNN1G*), distal convoluted tubules (*SLC12A3*), loop of Henle cells, thick ascending limbs (*SLC12A1*) and descending thin limbs (*SLC14A11*), podocytes (*NPHS2*), type A intercalated cells (*KIT*), type B intercalated cells (*SLC26A4*), principal cells (*AQP2*), mesangial cells (*PDGFRB*), endothelial cells (*PECAM1*), fibroblasts (*FBN1*), and immune cells (*CD36*, *IL7R*, *CD74*) (Fig. 1C-G). Comparison of our cell-type annotations with automated annotations, using single-cell kidney reference databases from humans (Azimuth) (Fig. 1H-I) (*42*) (*56*) and mouse (Fig. S3A-B) (*54*), showed high similarity. We also annotated a “proximal tubules-like” cell cluster, as this cluster expresses the proximal tubule markers *MIOX*, *SLC34A1*, and *LRP2* but at lower levels compared to the proximal tubules cluster (Fig. S3C). To confirm consistency of cell-type annotations before and after cross-species integration, we compared the annotations determined separately for each species before integration and the annotations determined jointly after integration and observed high similarity (Fig. 1J).

We detected several cell composition differences between frugivorous and insectivorous bat kidneys. Between the two bats, the insectivorous bat renal epithelial cell distribution most closely resembles that of the mouse (Fig. S3D) (*57*). In fruit bats, we found fewer thick ascending limbs (TAL) and significantly less descending thin limbs (DTL) cell-types (Fig. 1G), which make up the loop of Henle and are largely found in the renal medulla. The loop of Henle cluster was correlated with the insectivorous bat by CNA (Fig. S3E). Although TAL was not significantly more abundant in the insectivorous bat by Wilcoxon rank-sum test, likely due to small sample size, a Chi-square test of independence indicates high confidence of TAL enrichment in insectivorous bats (Fig. S3F). This is in line with previous reports that showed that fruit bats have a larger renal cortex and a smaller renal medulla compared to insectivorous bats (*16, 17, 58*). Additionally, we observed fruit bats to have significantly more type A intercalated cells, which are involved in acid secretion into the urine, and type B intercalated cells (Fig. 1G, Fig. S3F), which mediate bicarbonate secretion while reabsorbing sodium chloride (*59*), fitting with the high amounts of bicarbonate and the low amounts of sodium in fruit, which has a negative risk of renal acid load (*60*). The fruit bat also has significantly more principal cells, which reabsorb sodium and excrete potassium, than the insectivorous bat (Fig. 1G, Fig. S3F), in line with fruit containing low sodium and high potassium levels. All of these collecting duct cell-type clusters correlate with the fruit bat in CNA (Fig. S3E). The fruit bat also has significantly more connecting tubules, which are largely found in the renal cortex, and this cluster correlates with the fruit bat by CNA (Fig. 1G, Fig. S3E-F). Connecting tubules together with late distal convoluted tubules and the cortical collecting duct (principal cells, type A and B intercalated cells) are often called the aldosterone-sensitive distal nephron (ASDN). Aldosterone increases sodium reabsorption and promotes potassium secretion in the final step of the renin-angiotensin-aldosterone system (RAAS) (*61*). As hyperkalemia (excessive potassium in blood) stimulates aldosterone release (*61*), the greater abundance of connecting tubules in fruit bats is in line with a high potassium diet. The insectivorous bat has significantly more proximal tubule-like cells as compared to the fruit bat, and this cluster also shows significant association with insectivorous bats (Fig. 1G, Fig. S3E-F). Decreased proximal tubule count is also observed in human diabetic kidneys (*62*), although proximal tubular growth occurs in early diabetic nephropathy (*63*). In summary, we find that frugivorous and insectivorous kidneys differ in many nephron components, particularly in proximal tubules-like cells, connecting tubules, the loop of Henle and the collecting duct.

To further validate these cell composition differences, we performed immunofluorescence on various cell-type specific markers on kidney tissue sections from adult big brown bats and Jamaican fruit bats (Table S4). For type B intercalated cells, we used an antibody against pendrin, encoded by *SLC26A4*, finding significantly more pendrin-expressing cells in fruit bats than in insectivorous bats (Fig. 2A-B). For TAL, we used an antibody against the Na^+^/K^+^/2Cl^-^ co-transporter NKCC2, encoded by *SLC12A1*, finding significantly less NKCC2-expressing cells in fruit bats than in insectivorous bats (Fig. 2C). For principal cells, we used an antibody against aquaporin 2, encoded by *AQP2*, finding significantly more aquaporin 2-expressing cells in fruit bats than in insectivorous bats (Fig. S4A-B). Combined, our immunofluorescence results validate the single-cell composition differences observed for these cell-types.

**Fig. 2:**
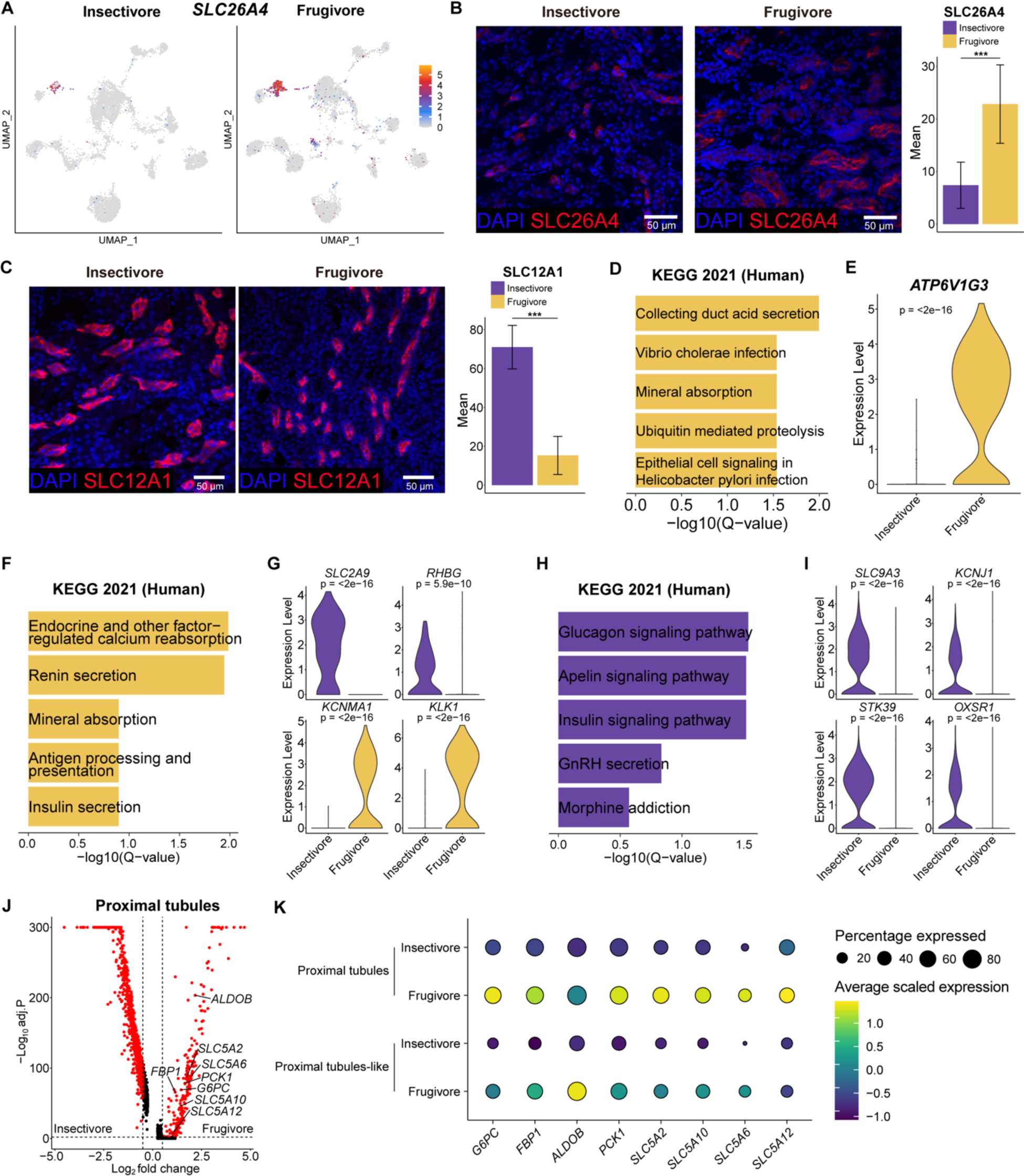
Immunofluorescence and differential gene expression analyses identify traits facilitating frugivory in the bat kidney. (**A**) UMAPs of type B intercalated cell marker gene *SLC26A4* expression in each species. (**B**) (Left) Representative images of *SLC26A4* immunofluorescence (red) in bat kidneys. Nuclei are stained with DAPI (blue). (Right) Quantification of *SLC26A4* immunofluorescence normalized to nuclei in bat kidneys. Results represent arbitrary units of fluorescence (AU) mean ± standard error of the mean (SEM) derived from 3 insectivorous and 3 frugivorous bats (n = 3/phenotype, n = 10 images/individual [see Materials and Methods]). Mixed effects model \*\*\**p*-value < .0001. (**C**) (Left) Representative images of *SLC12A1* immunofluorescence (red) in bat kidneys. Nuclei are stained with DAPI (blue). (Right) Quantification of *SLC12A1* immunofluorescence normalized to nuclei in bat kidneys. Results represent mean (AU) ± standard error of the mean (SEM) derived from 3 insectivorous and 3 frugivorous bats (n = 3/phenotype, n = 10 images/individual [see Materials and Methods]). Mixed effects model \*\*\**p*-value < .0001. (**D**) Bar plots showing Kyoto Encyclopedia of Genes and Genomes (KEGG) Human 2021 pathways enriched in fruit bat type B intercalated cells. (**E**) Violin plot of *ATP6V1G3* expression in type B intercalated cells. (**F**) Bar plots showing KEGG Human 2021 pathways enriched in fruit bat type A intercalated cells. (**G**) Violin plots of *SLC2A9*, *RHBG*, *KCNMA1*, and *KLK1* in type A intercalated cells. (**H**) Bar plots showing KEGG Human 2021 pathways enriched in insectivore thick ascending limbs. (**I**) Violin plots of *SLC9A3*, *OXSR1*, *STK39*, and *KCNJ1* expression in TAL. (**J**) Volcano plot showing differentially expressed genes between species in proximal tubules cells. (**K**) Dot plots showing the expression of gluconeogenesis and various SLC5 genes in bat proximal tubules and proximal tubules-like cells.

### Gene expression differences between insectivorous and frugivorous bat kidneys

We next analyzed our scRNA-seq datasets to identify molecular changes associated with adaptation to these dietary differences. We conducted gene ontology (GO) enrichment analyses for the genes that showed differential expression between frugivorous and insectivorous bat kidneys (Table S5-6). As there are significant cell-type composition differences in the kidney for collecting duct cells, we first focused on enriched terms identified in these cells. For type B intercalated cells, the top KEGG 2021 Human Pathway term for fruit bats was “collecting duct acid secretion”, involving the components of vacuolar H^+^-ATPase (Fig. 2D). For example, ATPase H^+^ transporting V1 subunit G3 (*ATP6V1G3*) was highly expressed in type B intercalated cells in fruit bats but not in insectivorous bats (Fig. 2E). Reabsorption of sodium in type B intercalated cells is fueled by H^+^-ATPase (*59*), so fruit bats most likely require more vacuolar H^+^-ATPase to reabsorb scarce sodium in their diet. For principal cells, the top GO Biological Process terms were all related to maintaining sodium-potassium balance (Fig. S4C-D), which includes fruit bat upregulated lysine-deficient protein kinases *WNK1* and *WNK4* (Fig. S4E). The WNK signaling pathway is regulated by sodium intake via aldosterone as well as insulin (*64*), and mutations in humans in both these genes lead to hyperkalemia and hypertension (*65*). Combined, our single-cell RNA-seq analyses reveal important gene expression adaptations to high dietary glucose and potassium and low dietary sodium and calcium that occurred within the collecting duct, many of which are associated with causing human disease like hyperkalemia and hypertension.

For type A intercalated cells, the top KEGG 2021 Human Pathway for fruit bats were “endocrine and other factor-related calcium reabsorption” and “renin secretion” (Fig. 2F). Amongst these genes was kallikrein 1 (*KLK1*), an endocrine-responsive gene that is induced by high glucose (*66*), and was found to be highly expressed in fruit bat type A intercalated cells but not in insectivorous bats (Fig. 2G). Interestingly, mutations in this gene or its promoter have been found to be associated with hypertension in humans, and its overexpression in mice leads to hypotension, protection from diabetic cardiac damage, renal fibrosis and renal vasodilation, and increased nitrate levels in urine (*67*), which are found naturally in high concentrations in plants (*68*). Furthermore, *KLK1* protects against hyperkalemia after a high potassium load in mice (*69*). Another highly expressed gene in the fruit bat type A intercalated cells that was not expressed in the insectivorous bat, was the potassium calcium-activated channel subfamily M alpha 1 (*KCNMA1*) (Fig. 2G). This gene is responsive to renin-mediated regulation of body potassium levels and localizes to the apical membrane of these cells under a high potassium diet to secrete potassium (*59*). Furthermore, high potassium stimulates the synthesis and function of the *KCNMA1*-encoded transporter (*59*), fitting the lack of expression in insectivorous bats. Increased expression of *KLK1*, *KCNMA1*, and vacuolar H^+^-ATPase in type A intercalated cells in fruit bats is likely an adaptation to fruit specialization that helps to address high glucose and high potassium intake.

To determine if TAL gene expression differed substantially between bat species, we used the STRING database (*70*) to identify genes related to *SLC12A1* and analyzed their expression in our datasets. We found that *STK39* (SPAK), *OXSR1* (OSR1), *KCNJ1* (ROMK), and *SLC9A3* (NHE3), key genes supporting and regulating TAL function in urine concentration (*71, 72*), showed significantly lower expression in fruit bats (Fig. 2H-J). *Stk39* null mice, as well as double knockout of *Stk39* and *Oxsr1*, have increased potassium in the urine, in line with the higher potassium levels observed in fruit bats (*73*). Mouse kidney tubule *Oxsr1* knockout, *Kcnj1* knockout, and *Slc9a3* conditional TAL knockout all show lower urine osmolality (*74–76*), fitting with their observed lower gene expression and the dilute urine found in fruit bats (*30*). Loss of function mutations in *SLC12A1* or *KCNJ1* in humans causes Bartter syndrome, which is characterized by hypokalemia, metabolic alkalosis (body pH elevation), polyuria (excess urination), salt wasting, hypercalciuria, and hypotension (*77–79*). Combined, our single-cell analyses find that both TAL abundance and gene expression changed substantially in fruit bats as compared to insectivorous bats.

We also analyzed our scRNA-seq data for expression of kidney transporters *SLC22A6* (OAT1), *SLC22A12* (URAT1), *SLC2A9* (GLUT9), and *RHBG*, which were reported to be lost in OWFBs (*20, 21*) but intact in NWFBs (Jamaican fruit bat and Honduran yellow-shouldered fruit bat, *Sturnira hondurensis*) (*21*), and for *MYO6* (myosin VI), which is highly expressed in the OWFB kidney (*80*). We found that *SLC22A6*, which is involved in the excretion of organic anions, and *SLC22A12,* which is involved in renal reabsorption of urate, show significantly higher expression in fruit bat proximal tubules and proximal tubule-like cells compared to those of the insectivorous bat (Fig. S4E). *SLC22A6* has extremely broad substrate specificity, for both endogenous and exogenous substrates (*81*), so in NWFBs it could possibly still have a role in the excretion of toxins. Higher expression of *SLC22A12* might be necessary to maintain urate homeostasis in fruit bats, as urate comes from endogenous purines and from diet (*82*), and fruits are low in purines (*83*). *SLC2A9*, which also mediates urate reabsorption in the kidney, was found to be highly expressed in type A intercalated cells in the insectivorous bat and showed no expression in the frugivorous bat in these cells (Fig. 2G). Lack of *SLC2A9* expression in the collecting duct cells in fruit bats could support urate reabsorption from a high glucose diet, as high glucose concentration inhibits urate reabsorption by this transporter (*84*). *RHBG*, which encodes an ammonium transporter, was expressed in type A intercalated cells in insectivorous bats but largely absent in fruit bats (Fig. 2G). Ammonium inhibits sodium reabsorption and potassium secretion (*85*), so reduced expression of *RHBG* would accommodate the low sodium-high potassium dietary intake of a frugivore. *MYO6* was upregulated in fruit bat podocytes (Fig. S4E). Adaptive evolution of *MYO6* in OWFBs was hypothesized to support protein preservation from a protein-scarce diet (*80*), as mice lacking *MYO6* had reduced endocytosis-mediated protein absorption in proximal tubules and also hypertension (*86*), which may similarly be supported by upregulation of *MYO6* in NWFB kidney. Together, these results suggest that kidney adaptations are not entirely synonymous across fruit bat lineages.

Bats have been evaluated as a model for diabetes due to their range in lifestyles, diet, and genetic factors (*3*). Features of diabetic kidneys include increased gluconeogenesis and glucose reabsorption (*87*). We scanned our datasets for gene expression signatures for these processes. We found gluconeogenesis genes *G6PC*, *FBP1*, *ALDOB*, and *PCK1* (*87*) to all be upregulated in fruit bat proximal tubules and proximal tubule-like cells, as well as the glucose reabsorption transporter *SLC5A2* (SGLT2) (*88*), along with other sodium/glucose cotransport family SLC5 genes *SLC5A10* (SGLT5), *SLC5A6*, and *SLC5A12* (Fig. 2J-K). Increased potassium secretion is also a signature of diabetic nephropathy (*62*), and we observed many gene expression changes in fruit bat kidneys that promote potassium secretion (*KCNMA1*, *STK39*, *OSR1*, *RHBG*) (Fig. 2G,I). Together, our gene expression data indicates that the fruit bat kidney shares many characteristics of a diabetic kidney.

### Gene regulatory differences between insectivorous and frugivorous bat kidneys

To analyze our scATAC-seq datasets and identify gene regulatory differences between insectivorous and frugivorous bat kidneys, we used MACS2 (*89*) to call 209,895 and 188,845 cell-type-specific peaks for these bat kidneys, respectively (Table S7-8). To associate open chromatin regions to genes and human phenotypes, we converted our bat peaks to human genome coordinates (*hg38*) (*90*), converting 141,779 insectivorous and 124,845 frugivorous bat peaks to *hg38*, respectively. We then used the Genomic Regions Enrichment of Annotations Tool (GREAT) (*91*) to characterize the target genes associated with these peaks. The fruit bat kidney was found to be highly enriched for human kidney phenotypes, including “glucose intolerance”, “metabolic alkalosis”, “abnormality of renin-angiotensin system”, “abnormal circulating renin”, and “hypokalemia” (Fig. S4F), fitting with our cell-type and scRNA-seq analyses that showed that the fruit bat kidney resembles a diabetic kidney. The insectivorous bat kidney was only enriched for the human kidney phenotype “enuresis nocturna” (involuntary urination) (Fig. S4F).

Next, to investigate differences in *cis*-regulation between frugivorous and insectivorous bats, we performed differential motif enrichment analyses on renal epithelial cells using AME (*92*). For collecting duct cells, we found two clusters of differentially enriched motifs, which separated frugivorous and insectivorous bats (Fig. 3A). Insectivorous bats were found to be enriched for *NKX* and *NFAT* motifs, and frugivorous bats were enriched for *ONECUT* motifs. *NKX* TFs are broadly expressed and important for organ development, and *NFAT* TFs control proteinuria (*93*), fitting with the high protein diet of insectivores. *NFAT5* upregulates ubiquitin ligase under hypertonic conditions in the collecting duct to protect renal medullary cells from apoptosis (*94*), which is also fitting for the low water insectivores obtain from their diet, as compared to frugivores, and the upregulation of *NFAT5* in insectivore principal cells and type A intercalated cells (Fig. S4G). *ONECUT1* is known to antagonize glucocorticoid-stimulated gene expression (*95*), regulate glucose metabolism (*96*), and mutations in this gene are associated with diabetes in humans (*97, 98*). Glucocorticoids stimulate expression of gluconeogenesis genes in the diabetic kidney (*62, 87*), fitting with the observed *ONECUT* motif enrichment which could allow tighter regulation of gluconeogenesis in response to fluctuating high glucose intake in frugivores. We also separated proximal tubules, proximal tubule-like cells, distal convoluted tubules, TAL and DTL for differential motif enrichment and found multiple TFBS clusters that differentiate frugivorous and insectivorous bats (Fig. 3A). The *NFAT5* motif was again enriched in insectivore tubules, and we also saw upregulation of *NFAT5* in insectivore proximal tubules and distal convoluted tubules (Fig. S4G). We found the *KLF9* motif to be enriched in fruit bat renal tubules (Fig. 3A), which was also enriched in diabetic proximal tubules (*87*). *KLF9* is a potential target for the mineralocorticoid receptor in mouse distal convoluted tubules (*99*), which mediates fluid and electrolyte balance (*100*) and undergoes non-ligand activation with elevated glucose (*101*). The *KLF9* motif was also enriched in fruit bat collecting duct cells (Fig. 3A). In summary, kidney TFBS differential enrichment between insectivorous and frugivorous bats identified key TFs involved in diet, with frugivorous bats demonstrating diabetic-associated motif signatures. We next sought to utilize our integrative single-cell datasets to survey diabetes-associated regions in bat kidneys. Examination of *KLK1*, which regulates blood pressure, showed a substantial increase in chromatin accessibility in the collecting duct cells as well as proximal tubules, TAL, and podocytes in the fruit bat (Fig. 3B). *ATP6V1G3*, which is a biomarker for diabetic nephropathy (*102*), showed greater chromatin accessibility, particularly in the promoter, in fruit bat type A and type B intercalated cells (Fig. S4H). The insectivore *ATP6V1G3* promoter is predicted to bind *RREB1*, which represses RAAS through the angiotensin gene and is associated with type 2 diabetes (T2D) end-stage kidney disease (*103*). Angiotensin is known to stimulate vacuolar H^+^-ATPase activity in intercalated cells (*104*), so the lack of the *RREB1* motif in the fruit bat *ATP6V1G3* promoter may allow greater activation of RAAS effects. In addition, *RREB1* expression is lower in fruit bat proximal tubules and proximal tubules-like cells (Fig. S4G). We also observed similar chromatin accessibility trends between human diabetic proximal tubules and controls (*87*) in bats (Fig. S5A-D). For example, *PCK1* demonstrated decreased accessibility near the end of the gene body in diabetic proximal tubules (*87*), which was also observed in the fruit bat kidney (Fig. S5A). The fruit bat sequence homologous to the insectivorous bat peak following the end of the *PCK1* gene body is predicted to bind transcriptional repressor *ZNF331* (*105*), fitting with the decreased accessibility of this region in fruit bat proximal tubules. Together, our analyses indicate that fruit bats evolved many features similar to a human diabetic kidney.

**Fig. 3:**
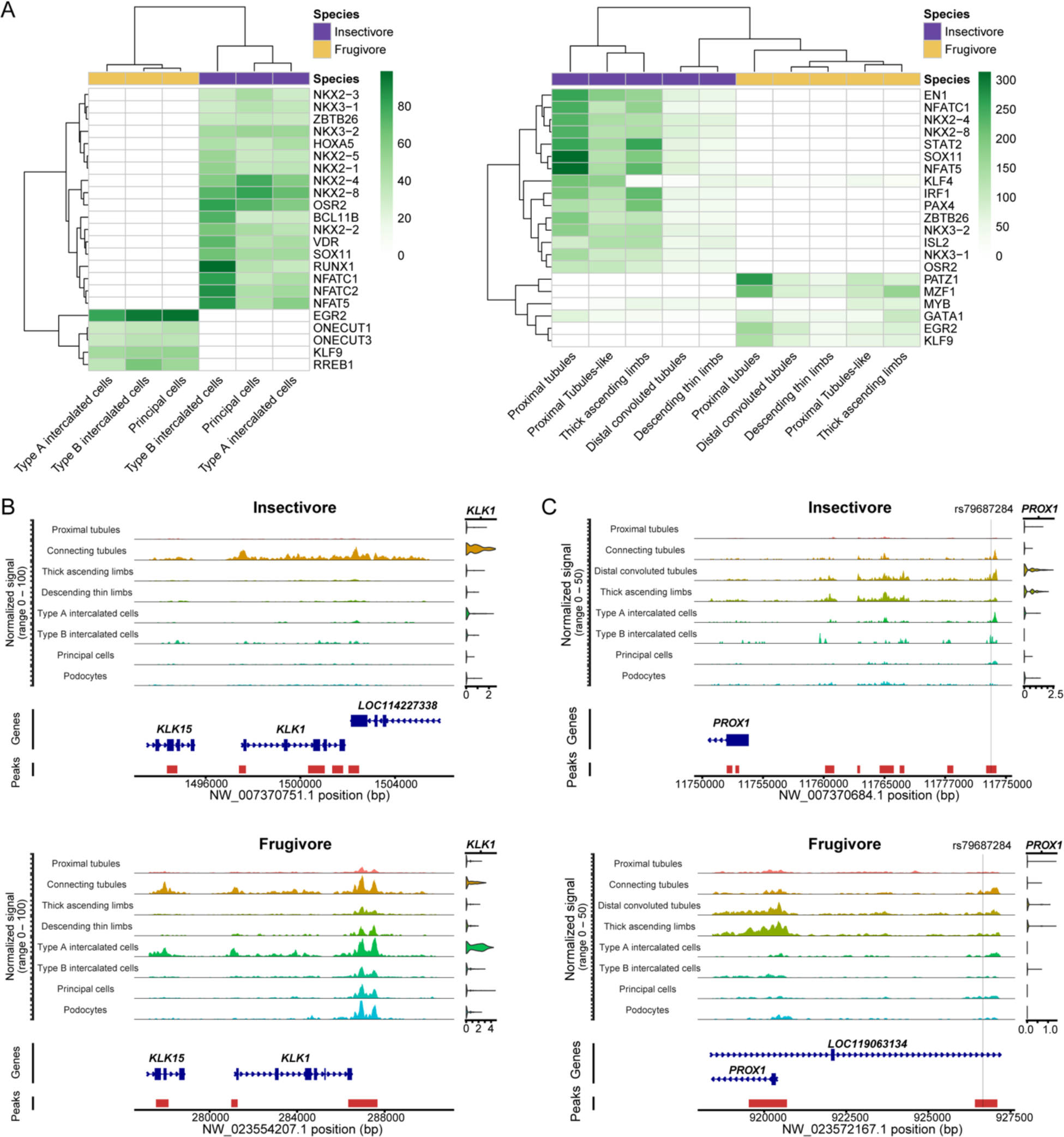
scATAC analyses reveal TFBS and chromatin accessibility in bat kidneys. (**A**) (Left) Heatmap of TF motifs enriched in bat kidney collecting duct cells. (Right) Heatmap of TF motifs enriched in bat renal tubule and limb cells. Both heatmaps display -log(p-value). (**B**) scATAC-seq coverage plots of *KLK1* in bats. SCTransform-normalized expression plot visualized on the right by cell-type. (**C**) scATAC-seq coverage plots of rs79687284 in the *PROX1* locus in bats. SCTransform-normalized expression plot visualized on the right by cell-type.

We next compiled a list of noncoding SNPs associated with type 1 diabetes (T1D) and T2D from the NHGRI-EBI Genome-Wide Association Study (GWAS) catalog (*106*). We then filtered this list with SNiPA (*107*) for noncoding single nucleotide polymorphisms (SNPs) that are in linkage disequilibrium (LD) with nearby variants (r^2^ > 0.8) (Table S9). We then determined the overlap of 3,460 noncoding SNPs with bat scATAC peaks in human genome coordinates, finding 381 overlaps (hypergeometric p-value < .0001) (Table S10), with the majority of these SNPs (∼90%) associated with T2D, including loci at *INSR* (*108*), *PIK3R1* (*109, 110*), and *MAP3K1* (*111–113*). Our cell-type-specific regulatory profilings allowed us to investigate human disease variants that are difficult to study without cell-type-specific data, such as rs79687284 at the *PROX1* locus.

This variant is highly associated with T2D and was predicted to have regulatory function as an enhancer, but it did not demonstrate transcriptional activity in luciferase reporter assays in a pancreatic beta cell line that expresses *PROX1* (*114*). We observed an increase in chromatin accessibility at this region in insectivorous bats and corresponding gene upregulation in insectivore distal convoluted tubules (Fig. 3C). *Prox1* heterozygous knockout (lower-expressing) mice have increased circulating insulin and glucagon and adult-onset obesity (*115*), and deletion of this gene in distal convoluted tubules causes hypomagnesemia (*116*), which is associated with progression of T2D (*117, 118*), and fitting with the magnesium-rich diet of insectivores (*119*). In summary, our scATAC-seq datasets allowed us to identify candidate gene regulatory elements in bats that overlap with T1D and T2D GWAS SNPs.

### Insectivorous and frugivorous bat pancreases have different cell-type composition

We next set out to analyze our multiome data for the bat pancreas. Similar to the kidney, we initially annotated all major cell-types using markers from previously reported scRNA-seq datasets from human and mouse pancreases (*53, 120, 121*). We detected all major known cell-types found in human and mouse pancreas: acinar cells (*CPA2*), ductal cells (*SLC4A4*), beta cells (*INS*), alpha cells (*GCG*), delta cells (*SST*), endothelial cells (*SLCO2A1*), active pancreatic stellate cells (aPSCs; *LAMA2*), quiescent pancreatic stellate cells (qPSCs; *COL3A1*), and immune cells (*ACTB*, *CD74*, *IL7R*, *MRC1*) (Fig. 4A-E). We identified gamma cells and epsilon cells by other known markers (*CHGA*; (*122*) and *ACSL1* (*121*), respectively), as many major markers were not shared across species. We observed high similarity between our cell-type annotations and automated annotations using human single-cell pancreas reference databases (Fig. 4F-G, Fig. S5A-B) (*42, 121*); (*120, 123–127*). Consistent with the adult human pancreas, acinar cells showed clear heterogeneity in the bat pancreas, with secretory acinar cells (acinar-s) expressing higher levels of digestive enzyme genes (*CPA1*, *PLA2G1B*, *SYCN*, *KLK1*, *CLPS*) and idling acinar cells (acinar-i) expressing lower levels of digestive enzyme genes (Fig. S6C). We compared the annotations determined separately for each species before integration and the annotations determined jointly after integration and observed high similarity (Fig. 4H).

**Fig. 4:**
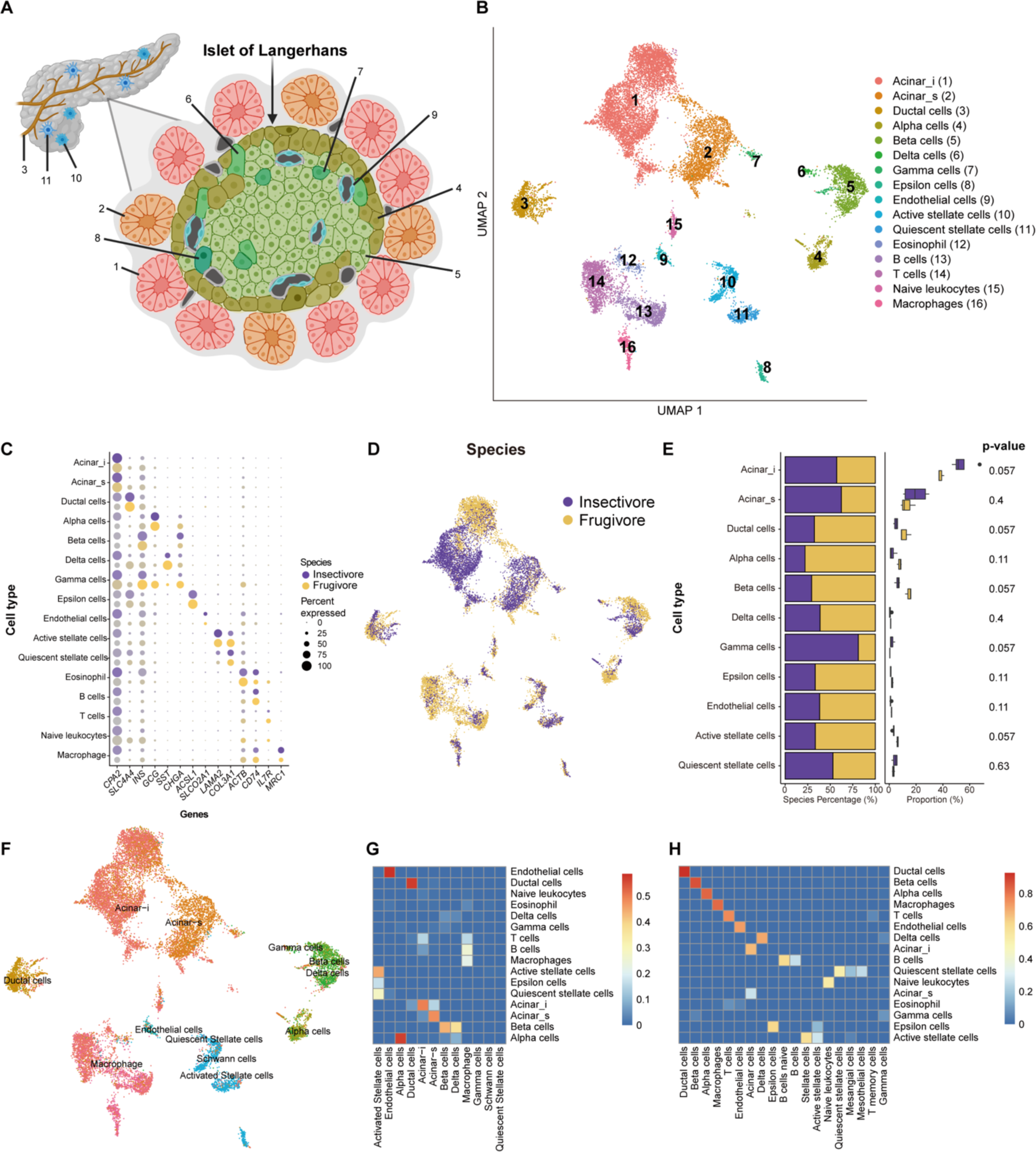
Joint scRNA and scATAC profiling of the bat pancreas. (**A**) Diagram of pancreas with zoom-in on the Islet of Langerhans. Colors correspond to cell-type colors in **B**. (**B**) UMAP of bat pancreas cell-types based on scRNA-seq data. (**C**) Dot plot of marker gene expression across all bat pancreas cell-types. Color intensity indicates the average expression level across all cells within a cell-type (purple or yellow is high, gray is low). (**D**) UMAP of bat pancreas cell-types by species/dietary phenotype. (**E**) Plot of species percentage across all pancreatic cells (left) with corresponding Wilcoxon rank-sum test for differential cell-type abundance (right). (**F**) UMAP of bat pancreas cell-types automatically annotated with human pancreas single-cell reference data (*121*). (**G**) Jaccard overlaps of auto-annotations in **G** (horizontal) with our integrated species annotations in **B** (vertical). (**H**) Jaccard overlap of all annotations found in each species before integration (horizontal) with after integration in **B** (vertical).

Fruit bats are reported to have an expansion of endocrine tissue in the pancreas (*18, 19*). Indeed, of the pancreatic cell-types, we found that the fruit bat pancreas is about 26% endocrine, whereas the insectivorous bat pancreas is about 12% (Fig. S6D). Both beta and alpha cell clusters are largely correlated with the fruit bat by CNA (Fig. S6E). As the smaller frugivorous bat sample size (N=3) precluded cell composition significance between species with Wilcoxon rank-sum test, we also conducted Chi-square tests of independence on Pearson residuals. We found, with high confidence, that both beta and alpha cells are more abundant in the fruit bat (Fig. S6F). Beta and alpha cells regulate blood glucose levels via insulin and glucagon, respectively, and their increased numbers are in line with the need to respond to a high glucose diet and tight blood glucose regulation in fruit bats, which have been shown to robustly regulate blood sugar in intraperitoneal glucose tolerance tests (*3, 22–24*). We found acinar cells to be more abundant in the insectivorous bat (Fig. S6E), with fruit bats having fewer acinar cells than insectivorous bats with high confidence (Fig. S6F). Acinar cells, which produce digestive enzymes for storage and secretion (*128*), comprise about 75% of insectivorous bat pancreatic cells, which is largely consistent with that of the human adult pancreas (*121*), compared to 52% in fruit bats (Fig. S6D). The substantial reduction in acinar cells in fruit bats could accommodate the increase of endocrine cells. The fruit bat also has more ductal cells with high confidence (Fig. S6F), which secrete enzymes from acinar cells into the duodenum and secrete bicarbonate to neutralize stomach acidity (*129*), although the overall percentage of ductal cells in the fruit bat pancreas (11%) corresponds with that of humans (10%) (*129*). In summary, our results suggest that compared to the insectivorous bat, the expanded endocrine pancreas in the fruit bat is largely attributable to increased numbers of beta and alpha cells, with proportionately lower exocrine acinar cell numbers that likely compensate for the endocrine expansion.

To validate our single-cell composition results, we performed immunofluorescence on pancreas tissue sections from the same bat species (Table S4). For beta cells, we used an antibody against insulin, encoded by *INS*, and found that there were significantly more insulin-expressing cells in fruit bats than in insectivorous bats (Fig. 5A). For alpha cells, we used an antibody against glucagon, encoded by *GCG*, and found that there were also significantly more glucagon-expressing cells in fruit bats than in insectivorous bats (Fig. 5B). Our immunofluorescence results thus validated our observed single-cell composition differences for endocrine cells.

**Fig. 5:**
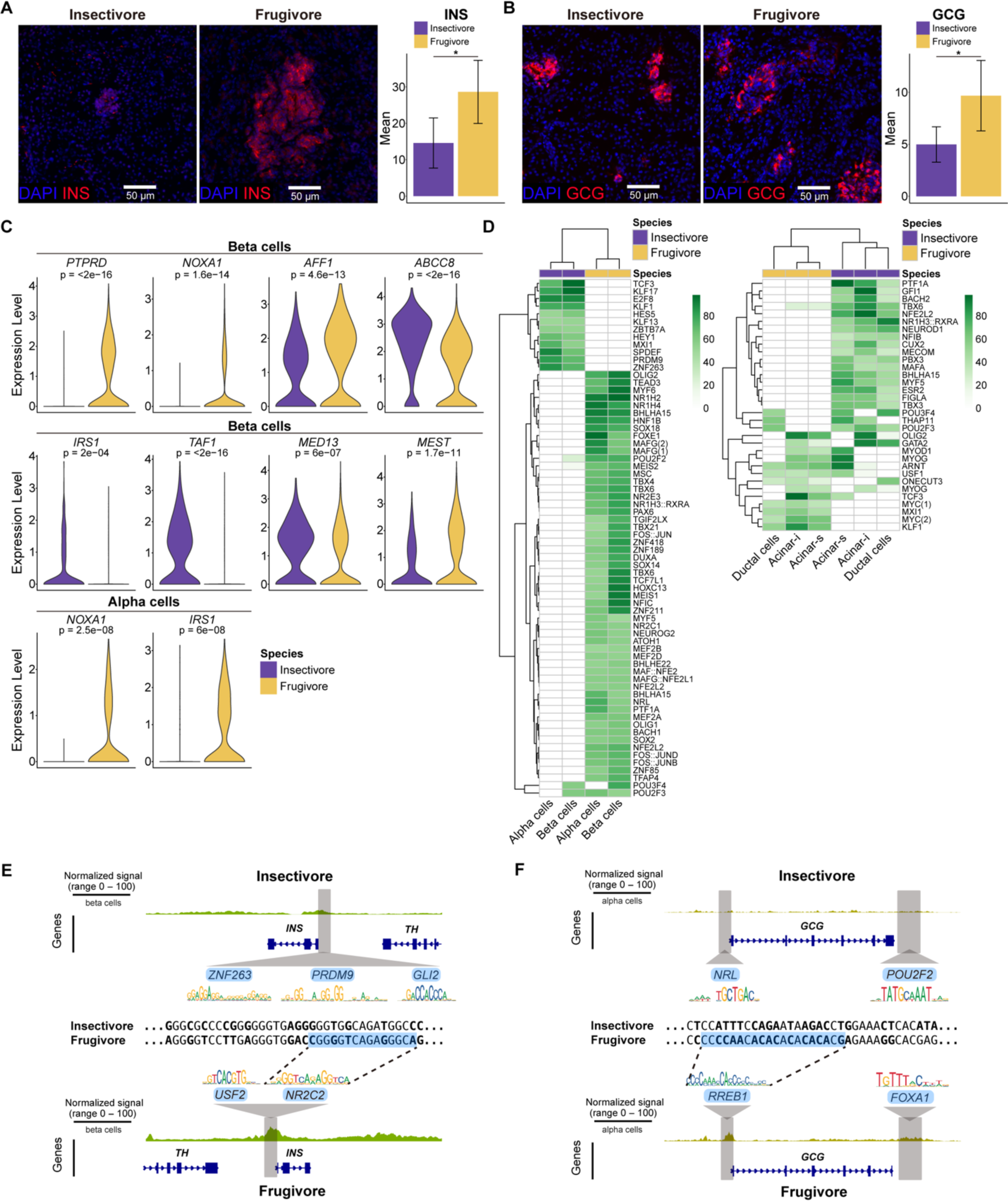
scRNA and scATAC analyses depict both exocrine and endocrine dietary differences between insectivorous and frugivorous bat pancreases. (**A**) (Left) Representative images of *INS* immunofluorescence (red) in bat pancreases. Nuclei are stained with DAPI (blue). (Right) Quantification of *INS* immunofluorescence normalized to nuclei in bat pancreases. Results represent arbitrary units of fluorescence (AU) mean ± standard error of the mean (SEM) derived from 3 insectivorous and 3 frugivorous bats (n = 3/phenotype, n = 10 images/individual [see Materials and Methods]). Mixed effects model \**p*-value = .002. (**B**) (Left) Representative images of *GCG* immunofluorescence (red) in bat pancreases. Nuclei are stained with DAPI (blue). (Right) Quantification of *GCG* immunofluorescence normalized to nuclei in bat pancreases. Results represent arbitrary units of fluorescence (AU) mean ± standard error of the mean (SEM) derived from 3 insectivorous and 3 frugivorous bats (n = 3/phenotype, n = 10 images/individual [see Materials and Methods]). Mixed effects model \**p*-value = .008. (**C**) Violin plots of diabetes-associated genes in bat endocrine cells. (**D**) (Left) Heatmap of TF motifs enriched in bat pancreatic endocrine cells. (Right) Heatmap of TF motifs enriched in bat pancreatic exocrine cells. Both heatmaps display -log(p-value). (**E**) scATAC-seq coverage plots of *INS* in bat pancreases with predicted TFBS highlighted in gray. Aligned sequence zoom-in on *NR2C2* motif with differing nucleotides depicted in bold. (**F**) scATAC-seq coverage plots of *GCG* in bat pancreases with predicted TFBS highlighted in gray. Aligned sequence zoom-in on *RREB1* motif with differing nucleotides depicted in bold.

### Gene expression differences between insectivorous and frugivorous bat pancreases

To identify pancreatic gene expression differences associated with frugivory, we performed gene enrichment analysis on all cell-types using differentially expressed genes between frugivorous and insectivorous bats (Table S11-12). As acinar-i cells appeared largely separated by species according to CNA (Fig. 4E, Fig. S6E), we did sub-clustering to analyze phenotypic differences within this cell-type (Fig. S7A), and found that cluster 0 (C0) was predominantly insectivorous bat-based and cluster 1 (C1) to be predominantly fruit bat-based. C0 was enriched for KEGG 2021 Human Pathway terms related to protein synthesis and secretion, fitting with the increased protein composition in insects. C1 was enriched for “diabetic cardiomyopathy” as well as “oxidative phosphorylation” (Fig. S7B-C), which is notably shared with human T2D acinar cells (*125*). Ductal cells, which produce alkaline-high pancreatic secretions and are regulated by calcium signaling (*130*), were enriched in fruit bats for “calcium signaling pathway” genes, including *PDE1A*, *EGFR*, and *ERBB4* (Fig. S7D-F). Fruit bats may stimulate more pancreatic secretions due to their low calcium diet to quickly digest carbohydrates and rapidly fuel their metabolism (*25*).

We next determined if genes suggested to be adaptive for frugivory in OWFBs were differentially expressed between our NWFBs and insectivorous bats. *FAM3B* and *FFAR3*, metabolic genes involved in insulin metabolism and signaling, were reported to be lost in OWFB genomes (*20, 21*). We observed both genes to be expressed in the NWFB pancreas (Fig. S7F). Expression of *FAM3B* was sparse throughout the NWFB pancreas as well as the insectivore pancreas. *FFAR3*, an inhibitor of insulin secretion (*131, 132*), was expressed in insectivorous bat beta cells and was nearly absent in fruit bats (Fig. S7F). Weak expression of *FFAR3* in NWFB pancreas is consistent with the hypothesis that loss of this gene in OWFBs is adaptive, as fruit bats secrete large amounts of insulin (*19*).

We then analyzed the expression of glucose transporters (GLUTs) in pancreatic cells. We first examined the main glucose transporter in beta cells (*133*), *SLC2A2* (GLUT2), which was previously shown to have a fruit bat-specific 11 bp deletion in its proximal promoter (*3*), and found that it was not differentially expressed between our insectivorous and frugivorous bats (Fig. S7F). Other GLUTs identified in human pancreatic cells also did not show differential expression between insectivorous and frugivorous bats, with the exception of *SLC2A13* (GLUT13), a H+/myo-inositol transporter found in both human endocrine and exocrine cells (*133*). *SLC2A13* showed significantly higher expression in fruit bat exocrine acinar-i and ductal cells (Fig. S7F). Myo-inositol is found in fruits like cantaloupe, which was consumed by our fruit bats, and is produced in tissues such as the kidneys (*134*). Increased sugar intake and diabetes increases the need for myo-inositol (*134*). Increased expression of *SLC2A13* in fruit bat exocrine pancreas complements the upregulation of myo-inositol oxygenase enzyme (*MIOX*), which catalyzes myo-inositol breakdown, in the liver of the same fruit bat species we analyzed (Jamaican fruit bat) but not in four insectivorous bat species (*135*). In line with these findings, we also observed upregulation of the myo-inositol transporter *SLC5A10* (*136*) in the fruit bat kidney proximal tubules (Fig. 2K).

Features of a diabetic pancreas include a reduced exocrine pancreas and loss of beta cells, with differences in the presence of insulitis and islet amyloidosis between T1D and T2D (*137*). We compared our gene expression data to reported single-cell datasets from the diabetic human pancreas (*125, 126, 138*). Differentially expressed genes between human T2D beta cells and normal beta cells, determined from three separate single-cell studies with a combined 19 T2D donors and 33 non-diabetic donors (*139*), included *TAF1*, *PTPRD*, *NOXA1*, *MED13*, *AFF1*, and *MEST*. These genes were also differentially expressed in our datasets between insectivorous and frugivorous bat beta cells (Fig. 5C). *PTPRD* is involved in insulin signaling and variants within this gene were found to be associated with gestational diabetes risk (*140, 141*). This gene was found to be upregulated in T2D beta cells and in our fruit bat beta, alpha, and ductal cells. *AFF1*, which has circular RNAs that control beta cell apoptosis (*142*), was also upregulated in both T2D and fruit bat beta and acinar cells. Chronic hyperglycemia is known to upregulate NAPDH oxidase (NOX) genes (*143*), and *NOXA1* (NOX activator 1) was found to be upregulated in our fruit bat beta, alpha, and acinar-i cells. We also observed the monogenic diabetes risk gene *ABCC8*, a regulator of potassium channels and insulin release, whose loss of function causes hyperinsulinism (excessive insulin secretion) and polyuria (*144*), to be downregulated in fruit bat beta cells. In addition, the insulin receptor substrate-1 (*IRS1*), whose deletion in mice causes hyperinsulinism but not diabetes (*145*), was found to be downregulated in fruit bat beta cells and upregulated in fruit bat alpha and acinar-i cells. Taken together, the fruit bat pancreas exhibits gene expression changes that are known to elevate and tightly regulate insulin secretion and signaling in response to a high sugar diet, many of which correspond to human diabetic gene expression dysregulation.

### Gene regulatory differences between insectivorous and frugivorous bat pancreases

To analyze open chromatin regions specific to cell-types in the pancreas, we called peaks with MACS2 (*89*), finding 187,421 and 281,823 cell-type-specific peaks in insectivorous and frugivorous bat pancreases, respectively (Table S13-14). To associate open chromatin regions to genes and human and mouse phenotypes, we identified 111,619 and 184,360 peaks from the insectivorous and frugivorous bat genomes, respectively, in humans (*hg38*). Analysis of these human peaks using GREAT (*91*) found the insectivorous bat pancreas to be enriched for many mouse phenotype terms involving fat metabolism, including “abnormal lipolysis”, “impaired lipolysis”, “increased inguinal fat pad weight”, “increased white fat cell number”, and “abnormal white fat cell number” (Fig. S7G), which could be indicative of adaptation to a higher fat diet from insects. The fruit bat pancreas was most enriched for human pancreas phenotype “metabolic alkalosis” (Fig. S7G), indicative of the high bicarbonate diet of frugivores.

We then examined differentially enriched TFBSs in scATAC-seq peaks between insectivorous and frugivorous bats in endocrine and exocrine cell-types. We found that there are two clusters of TFBSs that show differential enrichment across alpha and beta cells in the two bat genomes, resulting in separation of dietary phenotypes (Fig. 5D). Insectivorous bats were enriched for *TBX* and *MEF2* motifs, TFs with important roles in organ and embryonic development, respectively, and *HNF1B*, which controls endocrine precursor generation and when mutated is associated with maturity-onset diabetes of the young (MODY)(*146*). The enrichment for *HNF1B* motifs in insectivorous bats over fruit bats indicates differential regulation of beta cells in fruit bats. Fruit bats were enriched for *KLF* motifs. *KLF* TFs regulate functions and pathophysiology of the digestive system, many of which maintain beta cell function and control oxidative stress response genes (*147*). Two clusters of TFBSs were also annotated for exocrine cell-types, indicating again that there is a divergence in TFBS usage between these two bat species (Fig. 5D). Insectivore exocrine cells were enriched for *MAFA* and *NEUROD1* motifs, TFs that regulate insulin gene expression and beta cell development, respectively (*148*). *NEUROD1* was not found to be differentially expressed between bat exocrine cells, but was found to be upregulated in fruit bat alpha, beta, delta, and gamma cells (Fig. S7F). *NEUROD1* can switch cell fates from endocrine to exocrine in response to NOTCH signaling (*149*), so the lack of enrichment for *NEUROD1* in fruit bat exocrine cells may promote endocrine precursor cells and prevent exocrine cell development, consistent with the differences in cell abundances. Fruit bat ductal cells and acinar-i cells showed a motif enrichment for *USF1*, which is involved in the regulation of genes in response to high glucose including insulin (*150*) and the regulation of diabetic kidney disease (*151*), as well as for *ARNT*, which has reduced expression in diabetic human islets and its removal in mouse beta cells results in abnormal glucose tolerance and insulin secretion (*152*). Both *USF1* and *ARNT* were upregulated in fruit bat acinar-i cells (Fig. S7F). Taken together, fruit bats exhibit TFBS usage indicative of unique glucose and insulin regulation.

Next, we examined specific diabetes-associated regions for gene regulatory differences between insectivorous and frugivorous bats. We found more open chromatin in the promoter of *INS*, which was more highly expressed in fruit bat beta cells (Fig. 5E, Fig. S7F,H). In the promoter of *GCG*, as well as in a region downstream of *GCG*, we observed increased open chromatin in fruit bat alpha cells (Fig. 5F, Fig. S7I), although *GCG* was not differentially expressed between these species (Fig. S7F). The *INS* promoter in fruit bats is predicted to bind *NR2C2* and *USF2* (Fig. 5E). *NR2C2* is known to have a role in beta cell regulation and mice lacking *NR2C2* have hypoglycemia (*153*) and increased oxidative stress (*154*), while *USF2* expression is stimulated by high glucose and is known to control the synthesis of insulin (*150, 155, 156*). There were many high-scoring motif occurrences for the *INS* promoter in insectivorous bats, including *PRDM9*, which has a differentially enriched motif in fruit bat endocrine cells (Fig. 5D), *GLI2*, which can serve as a transcriptional repressor in embryonic development (*157*), and transcriptional repressor *ZNF263* (*158*) (Fig. 5E). The *GCG* promoter in fruit bats is predicted to bind *RREB1* (Fig. 5F). *RREB1* is known to potentiate the transcriptional activity of *NEUROD1*, fitting with the upregulation of this TF in fruit bat endocrine cells (Fig. S7F). The insectivore *GCG* promoter is predicted to bind *NRL* (Fig. S7I), a motif that was also identified in human *GCG* (*159*) and differentially enriched in insectivore endocrine cells (Fig. 5D). The fruit bat peak downstream of *GCG* is predicted to bind *FOXA1* (Fig. 5F), which regulates alpha cell differentiation, glucagon synthesis and secretion (*160*). The homologous sequence in insectivorous bats is predicted to bind *POU2F2* (Fig. 5F), a motif that was also differentially enriched in insectivore endocrine cells (Fig. 5D). The lack of differential expression of *GCG* between bats with different diets may be attributed to other *cis*- or *trans*-acting regulators. Of note, however, our scATAC-seq analyses found fruit bats to have both *GCG* and *INS* proximal and distal *cis*-regulatory regions that show a more open chromatin state compared to insectivorous bats (Fig. 5E-F, Fig. S7H-I).

We also intersected our T1D and T2D list of 3,460 noncoding SNPs, and overlapped them with bat pancreas scATAC peaks, and found 421 overlaps (hypergeometric p-value < .0001; >90% from T2D) (Table S15). These overlaps included variants at loci highly associated with human diabetes, including *FTO* (*161, 162*), *IGF2BP2* (*163, 164*), *CMIP* (*165, 166*), and *VPS13C* (*167, 168*). In addition, we are able to identify peak overlap with novel human diabetes variants like rs1947178, which is located 159,430 bp downstream of the *TOX* transcription start site (TSS) and is predicted to increase chromatin accessibility and gene expression of *TOX*, a transcriptional regulator in T1D (*169*). We observed increased accessibility in fruit bats in this region, along with upregulation of *TOX* in fruit bat beta, alpha, acinar-i and acinar-s cells (Fig. S8A). As *TOX* was also associated with diabetic nephropathy (*170*), we examined this locus in our kidney datasets and also observed increased accessibility at rs1947178 in fruit bat principal cells, combined with upregulation of *TOX* in these cells (Fig. S8B). In summary, our bat integrative single-cell datasets allowed us to identify gene regulatory elements and variants that are associated with diabetes in humans.

## Discussion

Using integrative single-cell sequencing, we characterized the cell populations, transcriptomes, and regulomes of insectivorous and frugivorous bat kidneys and pancreases in a single-cell manner. We identified major cell-types in the kidneys and pancreases of these bats, dissected the transcriptional and regulatory differences between them, and validated several cell composition findings with immunofluorescence. For the frugivore kidney, we found a reduction in loop of Henle cells, combined with a loss of urine-concentrating transporters. We observed an expansion of collecting duct cells, combined with upregulation of sodium reabsorption-potassium secretion genes, and genomic enrichment for disease phenotypes (Fig. 6). For the frugivore pancreas, we found an expansion of beta and alpha cells, accompanied by a reduction in acinar cells, and several genes and genomic regions associated with insulin secretion and signaling (Fig. 6).

**Fig. 6:**
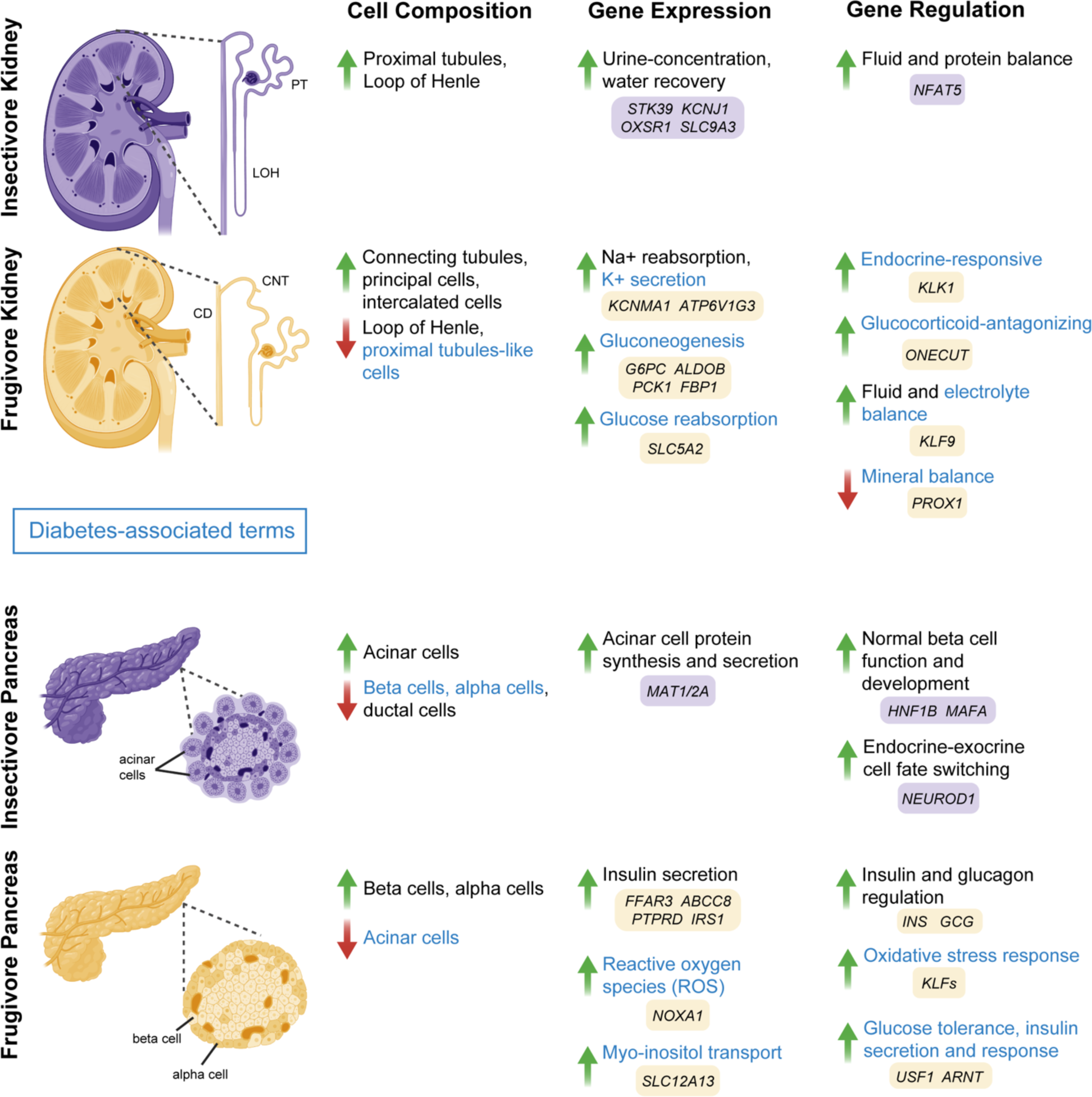
Summary of cell composition, gene expression and gene regulation differences between the bat insectivorous and frugivorous kidney and pancreas and how it relates to human diabetes. PT = proximal tubules, LOH = loop of Henle, CD = collecting duct, CNT = connecting tubules.

Combined, our work provides a cellular and molecular blueprint of frugivorous adaptation in mammals and can inform potential therapeutic targets for human disease, particularly hypertension, hyperkalemia and diabetes. To our knowledge, our results provide for the first time an unbiased and comprehensive analysis of the cell-type compositions that distinguish insectivorous and frugivorous mammals. For the kidney, we find that the medullary and cortical differences observed between frugivorous and insectivorous kidneys (*16, 17, 58*) are due to specific nephron composition differences in DTL, TAL, connecting tubules, principal cells, type A intercalated cells, type B intercalated cells, and proximal tubules-like cells (Fig. 6). As loop of Henle cells, DTL and TAL, are responsible for urine concentration and water recovery, and fruit bats get a substantial amount of water from fruit, fruit bats likely do not need as much of a structure for preserving water while excreting waste. A reduced renal medulla in response to water availability in diet or climate has also been observed in birds (*171–173*). The greater abundance of connecting tubules and collecting duct cells in fruit bats highlights nephron restructuring in response to high potassium and bicarbonate and low sodium and acid, favoring a larger ASDN. We also found less proximal tubules-like cells in fruit bats, which is notably observed in human diabetic kidneys (*62*). For the pancreas, we found that the large amount of endocrine tissue observed in fruit bats (*18, 19*) is due to greater beta and alpha cell abundances, relative to insectivorous bats, and a compensatory reduction in exocrine tissue, specifically in acinar cells (Fig. 6). These cell composition differences likely contribute to the unique ability of fruit bats to lower their blood sugar rapidly, even faster than insectivorous bats (*3, 22–24*). Taken together, our integrative single-cell analysis identified several novel cell composition differences and provides a cellular-level detailed catalog of previously observed morphological frugivorous kidney and pancreas features.

Prior to our study, identifying potential molecular adaptations to frugivory was mainly restricted to molecular evolutionary analyses of specific genes and comparative genomics (*3, 20, 21, 33*–*39, 80*). Here, we were able to use scRNA-seq to systematically identify gene expression differences in an unbiased manner. Our gene expression analyses identified numerous novel molecular adaptations that could be associated and vital for fruit specialization in mammals. In the kidney, the dilute urine observed in fruit bats (*30*) is likely attributed to decreased expression of key urine-concentrating transporters, *STK39*, *OXSR1*, *KCNJ1*, *SLC9A3*, within the TAL (Fig. 6). We also found that the fruit bat kidney exhibits gene expression changes to stimulate sodium reabsorption and potassium excretion, such as *KCNMA1*, *WNK1*, and *WNK4*, resembling an activated RAAS, which supports low sodium, high potassium dietary specialization (Fig. 6).

Some genes that were lost in OWFB genomes, like kidney transporters *SLC22A6* and *SLC22A12*, have been hypothesized to be adaptive for frugivory (*20*), and our data allowed us to determine whether these hypotheses are lineage-specific (loss of *SLC22A6* and *SLC22A12* is specific to OWFBs). In the pancreas, the increased sensitivity to insulin and glucose observed in fruit bats (*18, 19*) involves many genes that are associated with insulin secretion and signaling, like *PTPRD* and *ABCC8* (Fig. 6). Moreover, we discovered a unique connection in myo-inositol transport and metabolism from our data and previous gene expression analyses in the same fruit bat species (*135*) (Fig. 6).

Our scATAC-seq datasets allowed us to identify gene regulatory elements and TFBSs that could be involved in adaptation to frugivory. For the kidney, we found regulatory elements for many differentially expressed and diabetes-associated genes, including *KLK1*, which demonstrated cell-type-specific expression in bats with both promoter and enhancer accessibility, and *PCK1*, which was also shown to have differential chromatin accessibility downstream of the gene body in human diabetic proximal tubules (*87*). We also found insectivore renal epithelial cells to be enriched for *NFAT* motifs like *NFAT5*, which regulates proteinuria and osmotic pressure (*93, 94*), fitting the high protein consumption and relatively low water intake of insectivores, compared with that of frugivores. Frugivores were enriched for diabetes-associated motifs, such as *ONECUT*, and were highly enriched for human kidney phenotypes and for RAAS in particular. For the pancreas, we found *INS* and *GCG* promoters to be highly accessible in frugivore beta and alpha cells, respectively. As frugivorous bats are known to have remarkable blood sugar regulation (*3, 22–24*), promoter accessibility of *INS* and *GCG* may play a role. Insectivorous bats were also found to be enriched for motifs of many canonical pancreatic TFs like *HNF1B* and for many fat-related mouse phenotypes, corresponding to their higher fat diets, while frugivores were enriched for motifs of TFs related to diabetes like *KLF*s. As metabolites like lipids and carbohydrates regulate transcription through macronutrient-sensing TFs (*174*), the regulatory datasets we generated in this study offer the opportunity to investigate regulatory element evolution in response to dietary specialization.

The limitations of our study include our sample size, genome assembly and annotation, and the lack of functional genomic datasets for bats, precluding our ability to have known markers in bat kidneys and pancreases. Nonetheless, known markers in kidneys and pancreases of humans and mice were sufficient to identify major cell-types in the bat kidneys and pancreases. However, using human and mouse markers prevented us from identifying novel cell-types in bats. The smaller frugivore sample size for pancreas prevented us from detecting significant differences in cell composition, but we were still able to identify differences between bats with high confidence. Future studies may utilize spatial transcriptomics to compare cellular architecture and structures. We were also limited in analysis tools for non-model organism genomes. High-quality genomes are needed for deeper analyses, as they can provide better gene annotation, transcriptional isoforms and improve connection of gene regulatory elements to their target genes. Furthermore, novel genome annotations are needed to increase the number of shared features that can be recognized between species and to examine more genes in relation to humans and human disease. This was apparent in our results with the insectivorous bat genome having more annotated genes shared with humans than the fruit bat genome. Additional bat genomes, functional genomic databases, and more single-cell datasets from other insectivorous and frugivorous mammals will further determine whether our findings are specific to the species we investigated here, to NWFBs, and/or to other frugivorous mammals. Future studies with similar tissues could further investigate the differences between fasted and fed states within each bat species at various time points and understand developmental mechanisms for cell composition differences between the kidneys and pancreases of frugivorous and insectivorous bats. In addition, our study only focused on two organ types (kidney and pancreas) and thus could not uncover tissue adaptations in other organs that might be associated with frugivory.

Bats have been viewed as a model for diabetes research due to endocrine tissue differences between bat species and their unique blood sugar regulation. To the best of our knowledge, our study provides the first detailed analysis of the mammalian frugivorous kidney. The frugivore kidney exhibits many diabetic features, including decreased proximal tubules, upregulation of gluconeogenesis, glucose reabsorption, and potassium secretion genes, and corresponding diabetes-associated motifs, including *KLF9* which was also enriched in T2D renal tubules (*87*) (Fig. 6). Many of these kidney traits correspond with what we found in the frugivorous pancreas, such as increased transport and need for myo-inositol and motif and expression enrichment of diabetes-associated genes (Fig. 6). We also provide the first detailed analyses of bat pancreases, in which we documented differential expression of many signature genes in human T2D beta cells and differential enrichment of diabetes-associated motifs in endocrine and exocrine cells (Fig. 6). Together, our integrative single-cell study indicates that fruit bats evolved many diabetic-like features to deal with their diets but evolved protective mechanisms that prevent disease, such as upregulation of *KLK1* in type A intercalated cells in the kidney, which protects against diabetic tissue damage (*67*), and downregulation of *IRS1* in beta cells in the pancreas, which causes hyperinsulinism but not diabetes (*145*). Importantly, these data can be utilized to understand regulation of and adaptation to conditions that can cause disease in humans. We demonstrated how GWAS variants can be used with our single-cell data to reveal cell-type-specific regulation of diabetes-associated loci, such as *PROX1*, whereby rs79687284 was not previously examined in distal convoluted tubules or in the kidney. Our study also provides a unique perspective for human disease therapeutic development by revealing cellular and molecular adaptations to high sugar, potassium, and bicarbonate levels and to low amounts of protein, sodium, and calcium. *Cis*-regulation therapy (CRT) (*175*), for example, can take advantage of the genes and regulatory elements identified here for metabolic disease treatment. Moreover, our cell-type-specific data provides insight for designing localized treatments in heterogeneous tissues.

## Acknowledgements

The authors thank Carolyn Ku at UCSF for assistance with bat genome alignments. The authors also thank all the 2021 Belize “Batathon” members for assistance with bat collection. Fig. 1A-C, 4A, 6 and Fig. S1D were created with BioRender.com.

## Funding

National Human Genome Research Institute grant R01HG012396 (WEG, NA)

Korea Health Technology R&D Project, Korea Health Industry Development Institute (KHIDI),

Ministry of Health & Welfare, Republic of Korea grant HI19C1344 (SB)

California Institute for Regenerative Medicine (CIRM) Postdoctoral Fellowship (HPN)

UCSF Vision Core shared resource of the NIH/NEI P30 EY002162 (YMK)

UCSF CALM shared resource of the 1S10OD017993-01A1 (WEG)

National Institute on Deafness and Other Communication Disorders of the U.S. Public Health Service Research grant R01 DC00937 (AG)

Smithsonian National Museum of Natural History Peter Buck Postdoctoral Fellowship (MRI) National Institute of Allergy and Infectious Diseases (NIAID) grant R01AI134768 and R01AI140442 (TS)

National Science Foundation (NSF) grant 2020297257 (TS)

Evergrande Center startup funding (MH)

## Author Contributions

Conceptualization: WEG, NA

Methodology: WEG, SB, HPN, YMK, IGS, MH, NA

Fieldwork: WEG, MRI, NBS, NA Investigation: WEG, SB, HPN, RB, IGS Visualization: WEG, SB

Funding acquisition: WEG, SB, MH, NA

Project administration: AG, IL, MRI, NBS, TS, LNC, MH, NA Supervision: WEG, SB, MH, NA

Writing - original draft: WEG, SB, MH, NA

Writing - review & editing: WEG, SB, HPN, YMK, MRI, NBS, TS, LNC, IGS, MH, NA

### Data and materials availability

All data are available in the main text or the supplementary materials.

## Materials and Methods

### Bat samples and dietary treatment

For Chromium single-cell Multiome ATAC + Gene Expression experiments, kidneys and pancreases were obtained from four adult male *Artibeus jamaicensis* fruit-eating bats (Colorado State University [CSU]) and five adult male *Eptesicus fuscus* insect-eating bats (6-7 years old; Northeast Ohio Medical University [NEOMED]). All Jamaican fruit bats were fasted overnight during their night cycle (approximately 12 hours), and “fasted” bats were removed from their enclosures before feeding for euthanasia. “Fed” Jamaican fruit bats were provided unlimited cantaloupe and banana and were euthanized 30 minutes later. Tissues were harvested immediately for flash-freezing (May 2021) and stored at -80 Celsius until nuclei isolation. Because big brown bats feed on mealworms (*Tenebrio molitor*), we increased the fruit content of these mealworms to create a fruit “treatment” for insect-eating bats. These mealworms were fed a modified fruit diet for four days of sweet potatoes, apples, mango, and cane sugar. “Fasted” big brown bats were removed from their enclosures before feeding for euthanasia, and three “fed” big brown bats were fed an unlimited supply of fruit-enriched mealworms and were euthanized 30 minutes later. Tissues were harvested immediately for flash-freezing (all big brown bats were euthanized before the winter (September 2020) to exclude hibernation effects) and stored at -80 Celsius until nuclei isolation. All experimental procedures were approved under IACUC protocols (NEOMED IACUC #20-06-721, CSU IACUC #1034).

For immunofluorescence experiments, kidneys and pancreases were obtained from 3 adult male big brown bats of unknown age (June 2022; NEOMED IACUC #20-06-721), 2 adult male Jamaican fruit bats (August 2022; CSU IACUC #1034), and 1 adult female Jamaican fruit bat captured on an American Museum of Natural History (AMNH) field expedition to the Lamanai Archaeological Reserve in Orange Walk District of Belize in November 2021. The individual sampled (field number BZ701, voucher number AMNH.Mammalogy.281145) was caught in a ground-level mist net set near the High Temple at Lamanai (17.76750 N, 88.65207 W) on 17 November 2021. The bat was subjected to minimal handling after capture, and it was held in a clean cloth bag after capture as per best practices for field containment of bats. After species identification, the individual was euthanized humanely by isoflurane inhalation the same night it was captured. Capture and sampling were conducted under Belize Institute of Archaeology Permit IA/S/5/6/21(01) and Belize Forest Department Permit FD/WL/1/21(16), and samples were exported under Belize Forest Department permit FD/WL/7/22(08). All work was conducted with approval by the AMNH Institutional Animal Care and Use Committee (AMNH IACUC-20191212). Tissues were removed from the subject individual immediately following euthanasia and were flash-frozen in a liquid nitrogen dry shipper, with the cold chain maintained from field to museum to laboratory.

### Chromium single-cell multiome ATAC and gene expression sample processing and sequencing

For Chromium single-cell Multiome ATAC + Gene Expression experiments, we followed the manufacturer’s protocols for complex tissues (10x Genomics: CG000375, CG000338). Nuclei were sorted on a FACS Aria II (Becton Dickinson) using a capture target of 10,000 nuclei per sample to prepare libraries. ATAC libraries and GEX libraries were pooled together by tissue. Tissue-specific libraries were sequenced PE150 on two lanes of NovaSeq 6000 (Illumina) (table S1). The pancreas of a fed Jamaican fruit bat was not used due to low quality sequencing data.

### Bat genome modifications for joint scRNA and scATAC analysis

Jamaican fruit bat and big brown bat genomes and annotations were downloaded from NCBI (GenBank Assembly Accession: GCA_014825515.1, GCA_000308155.1). Scaffolds smaller than 50 kb for each genome were removed. Gene information in annotation files was collapsed so that each gene was represented by a single “exon” transcript. A mitochondrial genome was generated for the big brown bat using GetOrganelle (*44*), with default parameters for animal mitochondria assembly, and the output fasta was input into MITOS (*176*), with vertebrate genetic code, to create the corresponding gtf file (GenBank TPA: BK063052). Each species annotation was modified with one-to-one orthologue IDs created from Orthofinder (*45*) (table S16), which takes proteomes as input and assigned 89.9% of the total bat genes.

### Individual sample scRNA-seq and scATAC-seq processing and quality control (QC)

A total of 15 scRNA and scATAC FASTQs were input into cellranger-arc 2.0 (10x Genomics) whereby raw feature barcode matrices were generated with “count”. ATAC matrices were loaded as an object into Seurat version 4.0 (*42*). A common peak set was created within a species with R package GenomicRanges 1.50.2 (*46*) “reduce”. RNA matrices were then combined with ATAC matrices using only cells that overlapped the existing object. Cells were filtered for mitochondrial percentage < 25%; ATAC peak count 500 < x < 100,000; RNA count 200 < x < 25,000; nucleosome signal < 2; TSS enrichment >1 (terms defined by ENCODE (*177*)).

### Joint scRNA-seq and scATAC-seq bioinformatics workflow within species

Using post-QC Seurat objects as generated above, SCTransform was used to normalize each sample with mitochondrial percentage as a regressed variable and to find the top 3,000 variable features from each sample. Species replicates were then merged. To integrate RNA (SCT) across all samples within a species, 3,000 repeatedly variable features across each sample were selected with “SelectIntegrationFeatures”, and Harmony was run on the PCA dimensions 1:30, removing original sample identification as a variable. To integrate ATAC across all samples within a species, peaks were normalized with Signac function “RunTFIDF”, the most frequently observed features were identified with Signac version 1.8.0 (*46*) function “FindTopFeatures” with “min.cutoff = ‘q0’”, and Harmony was run on the singular value decomposition (SVD) dimensions 2:30, removing original sample identification as a variable. To integrate the integrated RNA and ATAC modalities, weighted nearest neighbor analysis (*42*) was applied on the Harmony reductions using 30 dimensions for each modality. Seurat function “FindClusters” was used with SLM algorithm and 0.6 resolution to identify clusters. Seurat function “FindAllMarkers” was used to identify cluster-specific markers and manually assign clusters according to shared markers to known mouse and human cell-types.

### Joint scRNA-seq and scATAC-seq bioinformatics workflow across species

Using post-QC Seurat objects mentioned above, gene activity matrices were created for each sample with the Seurat function “GeneActivity”, and ATAC peaks were removed from each Seurat object. R package SCTransform (*178*) was used to normalize each sample with mitochondrial percentage as a regressed variable and to find the top 3,000 variable features from each sample. Samples of the same species were then merged while retaining SCT matrices. Merged gene activity matrices were then log-normalized with the Seurat function “NormalizeData” with “scale.factor” set to the median of the gene activity counts. Two-thousand variable features were detected for normalized gene activity scores for each species with Seurat function “FindVariableFeatures” (vst method). Species were then merged. To integrate RNA (SCT) across species, we used Seurat function “SelectIntegrationFeatures” to select the 3000 top scoring variable features across all samples and manually added the selected variable features to the merged samples, and Harmony (*47*) was run on the principal component analysis (PCA) dimensions 1:30 with 20 maximum iterations and removing original sample and species variables. To integrate gene activity scores across species, we used Seurat function “SelectIntegrationFeatures” to select the 2,000 top scoring variable features across all samples and manually added the selected variable features to the merged samples. The features were scaled with Seurat function “ScaleData”, and Harmony was run on the PCA dimensions 1:30, removing original sample identification and species variables. To integrate the RNA and gene activity score modalities, weighted nearest neighbor analysis (*42*) was applied on the Harmony reductions using 30 dimensions for each modality. Seurat function “FindClusters” was used with SLM algorithm and 0.6 resolution to identify clusters, and “FindSubClusters” was used with original Louvain algorithm and 0.1 resolution to identify subclusters in pancreatic acinar cells. Seurat function “FindAllMarkers” was used to identify cluster-specific markers and manually assign clusters according to shared canonical markers to known mouse and human cell-types.

### Cell composition analyses

We calculated cell-type proportions from each sample and compared proportion percentages between the species by calculating significance with the Wilcoxon signed-rank test. The resulting boxplots for proportion percentages for each species and p-values were visualized with R package ggplot2 version 3.4.0 (*179*). For calculating species or condition associations per cell, we used R package CNA version 0.0.99 (*52*). For species associations, we used species variables as testing variables and used the neighbor graph generated from Seurat’s Weighted Nearest Neighbor Analysis from above. For condition associations, we used condition variables as testing variables and regressed out effects from species variables. For mosaic plots and calculation of p-values with pearson residuals, we used R package VCD version 1.4.10 (*180*) with species information and cell-type annotations.

### Automated cell-type annotations with human and mouse references and comparison

For the automated cell-type annotation method using human and mouse single-cell references, we used R package Azimuth version 0.4.6 (*42*) with built-in Azimuth human kidney dataset (*42*); (*56*), mouse p0 and adult kidney (*54*), Azimuth human pancreas dataset (*42, 121*); (*120, 123*–*127*), and human adult pancreas (*121*). For reference datasets with several levels of annotations, the most major cell-types were selected. For comparing similarity between manual annotations and automated annotations, we calculated Jaccard similarity coefficients between cells from each cell-type with different annotation methods. The resulting similarity heatmaps were visualized with R package pheatmap version 1.0.12 (*181*).

### Immunofluorescence staining, imaging and analysis

Flash-frozen samples were embedded in Tissue-Tek OCT Compound (Sakura Finetek, Torrance, Ca, USA), cut as 14uM cryo-sections, and fixed in 4% paraformaldehyde for 10 minutes at room temperature. Sections were rinsed in Phosphate Buffered Saline (PBS) and blocked in a humidity chamber in 3% normal donkey serum (Sigma-Aldrich, 566460) and 0.3% Triton X-100 (Sigma-Aldrich, T8787) in PBS. Sections were then incubated overnight at 4 degrees Celsius in a humidity chamber with one of the following protein-targeting primary antibodies from Thermo Scientific^TM^ at respective dilutions in blocking buffer: *SLC12A1* (18970-1-AP; 1:100), *AQP2* (PA5-78808; 1:100), *SLC26A4* (PA5-115911; 1:100), *INS* (15848-1-AP; 1:500), *GCG* (15954-1-AP; 1:500). Epitope matching can be found in table S4. Sections were washed with PBS and incubated in a humidity chamber for two hours at room temperature with secondary antibody Donkey Anti-Rabbit IgG NorthernLights^TM^ NL557-conjugated Antibody (Bio-techne, NL004; 1:200). Sections were rinsed with PBS and either incubated with Hoechst 33342 Solution (20mM) (Thermo Scientific^TM^, 62249; 1:10000) for 5 minutes before being mounted with ProLong^TM^ Diamond Antifade Mountant (Invitrogen^TM^, P36970) or mounted with ProLong^TM^ Diamond Antifade Mountant with DAPI (Invitrogen^TM^, P36962). Tissue sections were imaged on a CSU-W1 Spinning Disk/High Speed Widefield microscope with a Plan-Apochromat 20x objective and the Andor Zyla 4.2 sCMOS camera for confocal imaging or the Andor DU-888 EMCCD camera at the UCSF Center for Advanced Light Microscopy (CALM). On a section of tissue, ten random z-stack images (2-22 sections at .92uM thickness) were collected using the same imaging specifications respective to the target antibody. Fluorescence illumination was kept to a minimum to avoid photobleaching. All images shown are single z-sections processed with ImageJ on the FIJI platform (*182*) using the same processing parameters respective to the target antibody.

Tissue section nuclei count and total antibody fluorescence intensity were measured for every z-section for each image with CellProfiler (*183*) using the same detection framework respective to the organ and to the target antibody. A fluorescence intensity threshold was created for each antibody to reduce background signal. Nuclei counts and total antibody fluorescence intensities (arbitrary units[AU]) were summed across each image z-stack. The sum total antibody intensity was normalized to the sum nuclei count to obtain total antibody intensity (AU)/nuclei for each of the 10 images per tissue section. These 10 normalizations were then averaged to get the tissue section average of total antibody intensity (AU)/nuclei. To obtain the species average of total antibody intensity (AU)/nuclei, we averaged the tissue section averages by species. Results of immunofluorescence experiments represent mean ± standard error of the mean (SEM) derived from 3 individual insectivorous bats and 3 frugivorous bats (n = 3/phenotype). *P* values were calculated using a mixed effects model with restricted maximum likelihood and Satterthwaite approximation for degrees of freedom.

### Differential expression and gene set enrichment analyses

We used Seurat function “FindMarkers” for calculation of differentially expressed genes. We used a threshold of average log base2 fold change > 0.25, adjusted p-values < 0.01, and minimum detected gene fraction (min.pct) > 0.25. For DEGs between species, we additionally filtered out genes that are not expressed in both species. The differentially expressed genes were visualized with volcano plots using R package ggplot2 version 3.4.0 (*179*). We visualized selected genes on UMAP embeddings with Seurat function “FeaturePlot” and “VlnPlot” for violin plots. For gene set enrichment analysis using DEGs, we selected top genes with smaller counts of 100 or all DEG counts for each condition. We used R package EnrichR version 3.0 (*184*) for stated pathway databases and *p*-value calculations. The gene set enrichment results were visualized with ggplot2 with adjusted *p*-values calculated with EnrichR.

### Peak calling and enriched motif and GREAT analyses

ATAC data was added back into the integrated Seurat objects (fig. S1D) following separation by species via the Seurat function “SplitObject()”. Peaks were called for each cell-type by MACS2 (*89*) with “--nomodel --extsize 200 --shift -100” and effective genome size respective to each bat genome. Cell-type-specific peaks were associated genes and human and mouse phenotypes with the Genomic Regions Enrichment of Annotations Tool (GREAT; (*91*). GREAT only takes human or mouse coordinates for input, therefore big brown bat peaks were lifted to the human genome *hg38* (*90*) using UCSC Genome Bioinformatics Group tools chainSwap and liftOver (*185*) with -minMatch=.1 and chain file “hg38.eptFus1.all.chain”. To lift Jamaican fruit bat peaks to *hg38* for input into GREAT, a chain file was created between *hg38* chromosomes 1-22 and tandem repeat-masked Jamaican fruit bat genome via Lastz (*186*) and UCSC Genome Bioinformatics Group tools trfBig, mafToPsl and pslToChain (*185*). GREAT defines a basal regulatory domain for a gene as 5 kb upstream and 1 kb downstream of the gene’s TSS, regardless of other nearby genes, and extends this domain in both directions to the nearest gene’s basal domain, but no more than 1,000 kb in one direction. We viewed outputs from “Significant By Region-based Binomial” view.

Differentially enriched TFBSs from cell-type-specific peaks were estimated from AME software within the MEME suite (*187*), setting each time the other species’ cell-type-specific peaks as control sequences and default parameters. TFBS prediction on regulatory sequences was executed with FIMO software within the MEME suite (*187*) and filtered for *p*-value < .0000006 and *q*-value < .0003. For AME and FIMO motif inputs, we used the JASPAR 2022 core redundant dataset for vertebrates (*188*).

### Diabetes SNP curation and intersection with scATAC-seq peaks

SNPs associated with T1 and T2D were downloaded from the NHGRI-EBI GWAS Catalog (*106*). These SNPs were filtered for noncoding regions and proxied for LD with SNiPA (*107*) and threshold r^2^ > .8. The *hg38* coordinates for the resulting SNPs were intersected with the human genome coordinates of the bat cell-type-specific ATAC peaks. Bat scATAC-seq coverage over these human variants was examined by lifting the human genome coordinates back to bats using liftOver (*185*) and the chain files described in GREAT analyses. Hypergeometric p-values for SNP enrichment in bat ATAC peaks were calculated as follows: population size (N) = 611,564,276 noncoding SNPs from NCBI dbSNP 151 (*189*); number of success in the population (M) = 10,059 or 9,700 noncoding dbSNPs overlapping bat kidney or pancreas peaks, respectively; sample size (s) = 3,460 T1/2D SNPs; number of successes (k) = 381 or 421 T1/2D SNPs overlapping bat kidney peaks or pancreas peaks, respectively.

---

**Fig. S1.**
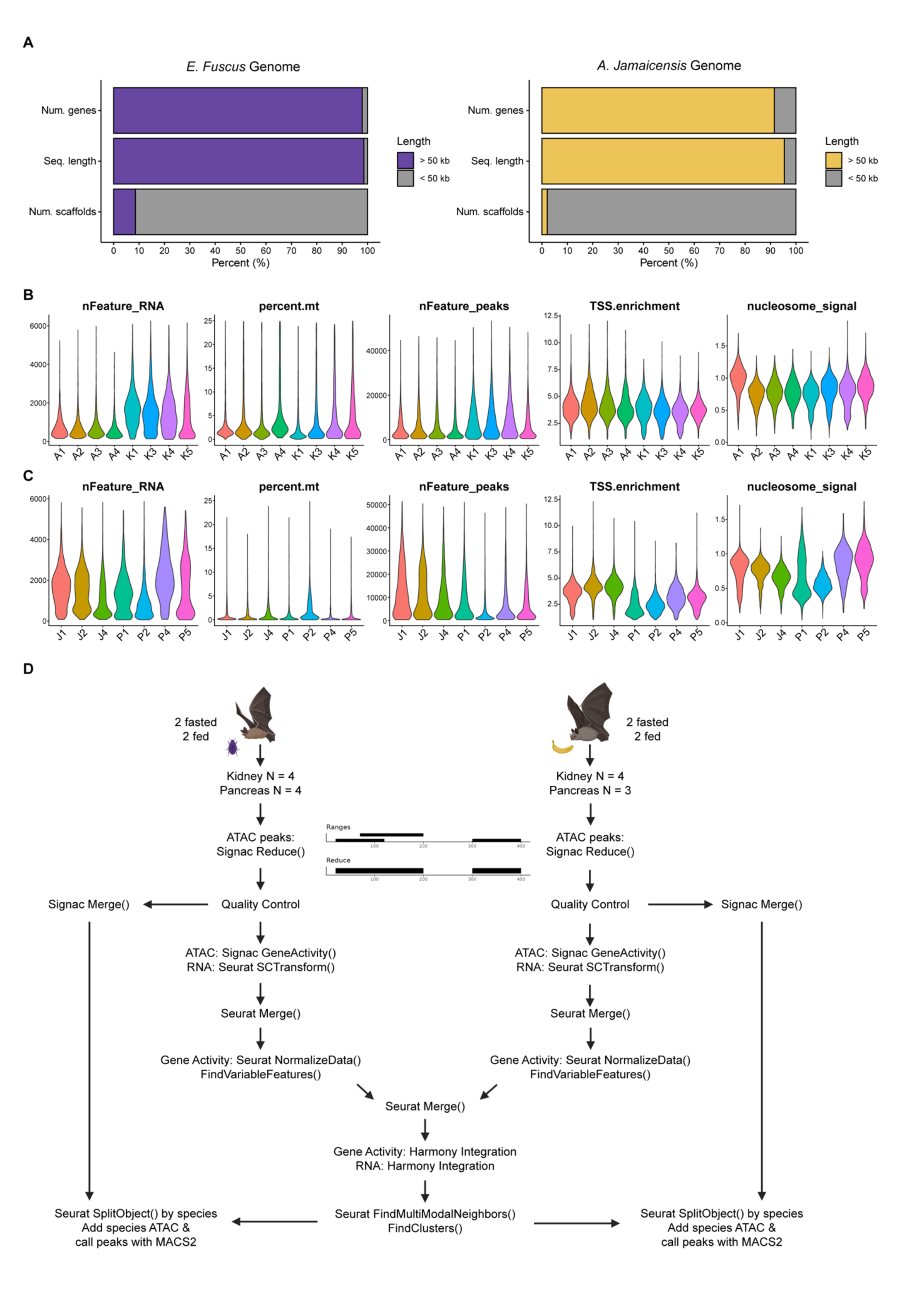
Joint scRNA and scATAC design and processing in bat tissues. (**A**) Bar chart showing that the number of genes and the genome sequence length captured by the number of scaffolds > 50 kb is > 90% for both bat genomes used in this study. (**B**) Violin plots of QC metrics Num. RNA features (nFeature_RNA), mitochondrial percentage (percent.mt), Num. ATAC features (nFeature_peaks), transcription start site enrichment (TSS.enrichment), and nucleosome signal for single-cell multiome on bat kidneys. A1-2 = fasted fruit bats, A3-4 = treated fruit bats, K1-3 = treated insect bats, K4-5 = fasted insect bats. (**C**) Violin plots of QC metrics nFeature_RNA, percent.mt, nFeature_peaks, TSS.enrichment, and nucleosome signal for single-cell multiome on bat pancreases. J1-2 = fasted fruit bats, J4= treated fruit bat, P1-2 = treated insect bats, P4-5 = fasted insect bats. (**D**) Schematic for cross species integration of multi-omic data (*46*).

**Fig. S2.**
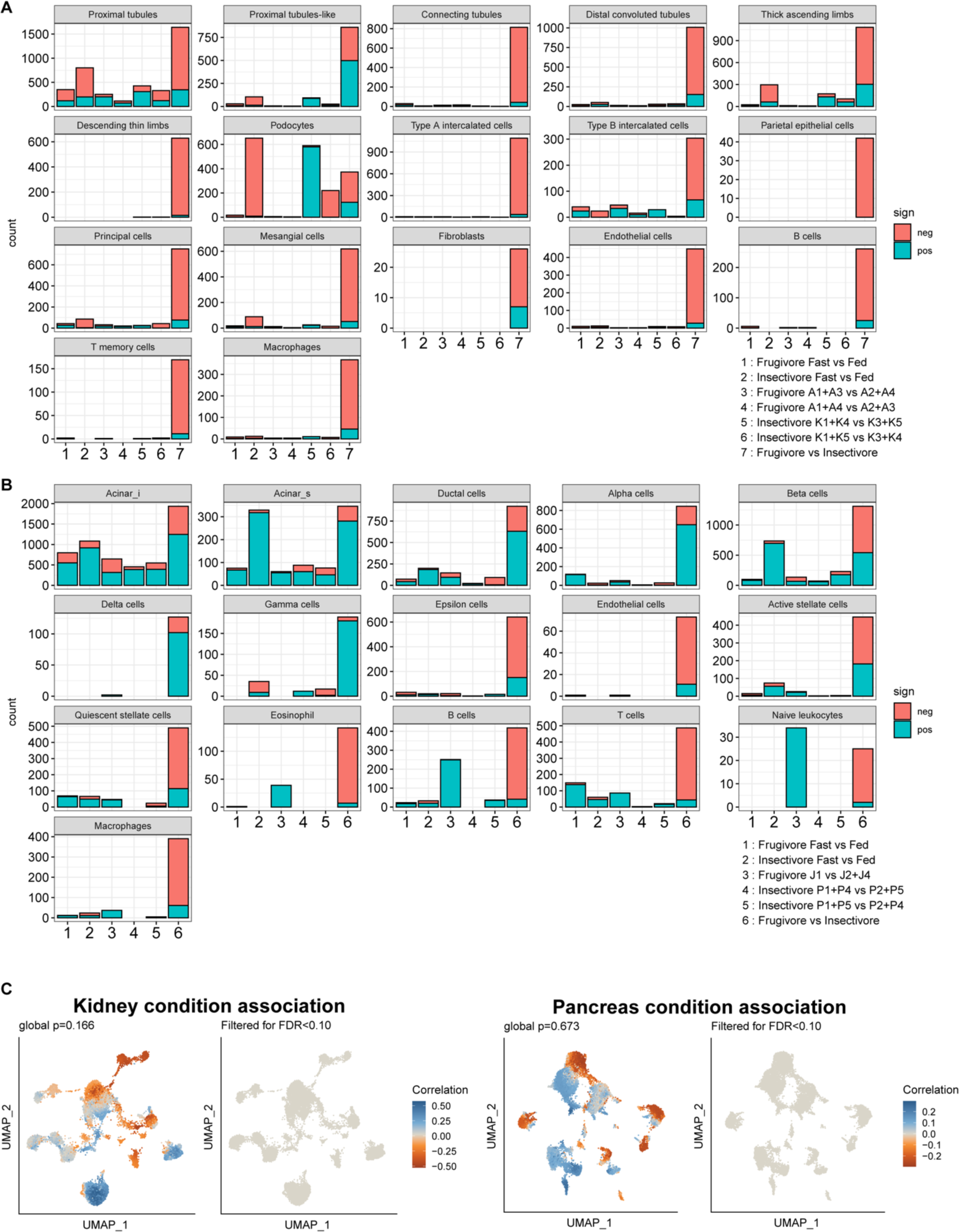
Differential expressed gene (DEG) counts by condition and by species in bat kidney and pancreas. (**A**) Bar charts of DEG counts in bat kidneys (legend notation: pos. vs neg.). A1-2 = fasted fruit bats, A3-4 = treated fruit bats, K1-3 = treated insect bats, K4-5 = fasted insect bats. (**B**) Bar charts of DEG counts in bat pancreases (legend notation: pos. vs neg.). J1-2 = fasted fruit bats, J4= treated fruit bat, P1-2 = treated insect bats, P4-5 = fasted insect bats. (**C**) (Left) CNA of condition associations to cell-types in bat kidney. (Right) CNA of condition associations to cell-types in bat pancreas.

**Fig. S3.**
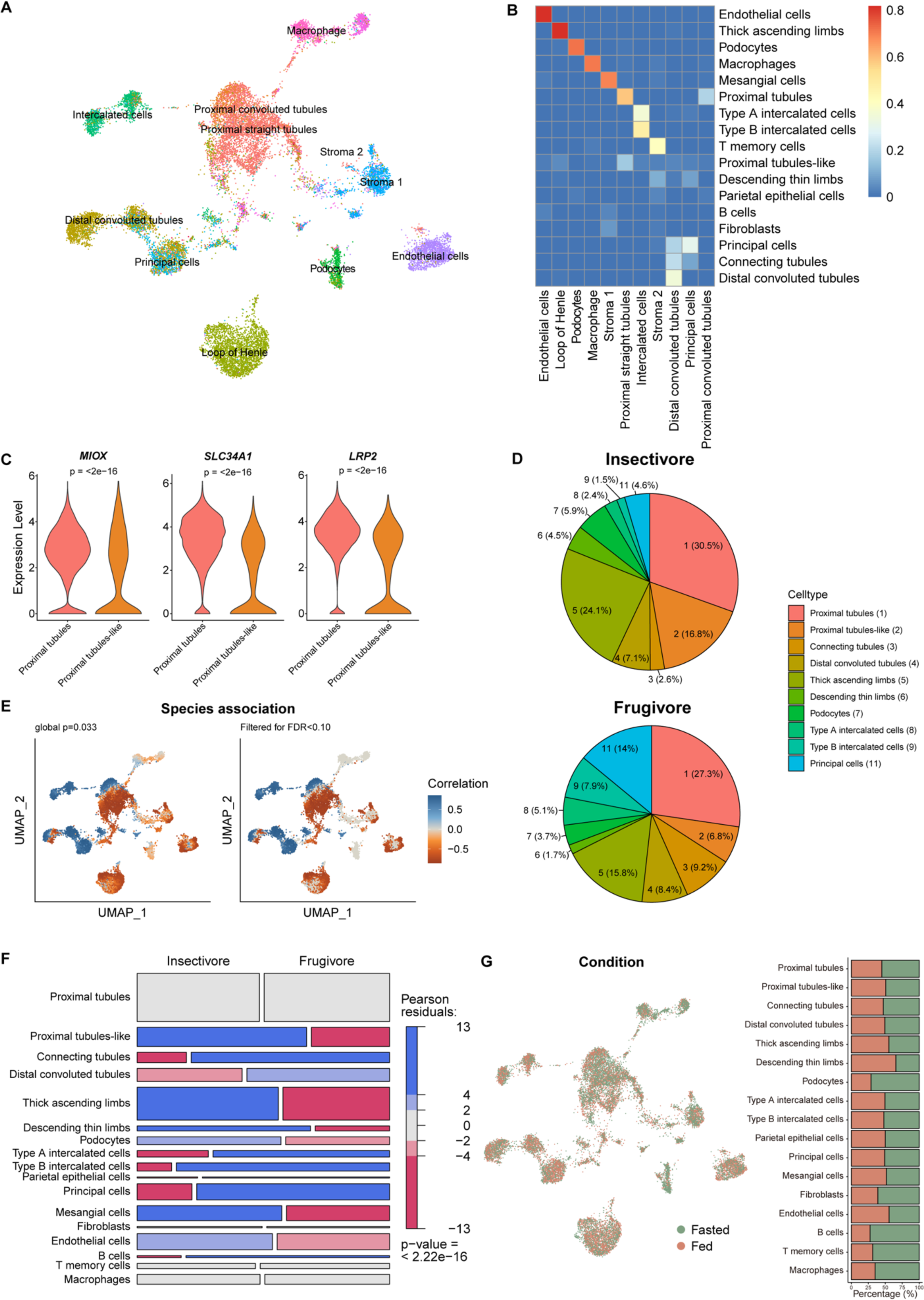
Single-cell composition analysis of bat kidney. (**A**) UMAP of bat kidney cell-types automatically annotated with mouse kidney single-cell reference data from Azimuth (*42*). (**B**) Jaccard overlaps of auto-annotations in **A** (horizontal) with our integrated species annotations in Fig. 1D (vertical). (**C**) Violin plots of proximal tubule marker gene expression in bat proximal tubules and proximal tubules-like cells. (**D**) Pie charts of cell-type percentages across renal epithelial cells. (**E**) CNA of species associations to cell-types (**F**) Pearson residuals visualized by mosaic plot of all cell-types identified in bat kidneys. (**G**) UMAP of bat kidney cell-types by condition. The bar chart shows the proportion of condition for each cell-type.

**Fig. S4.**
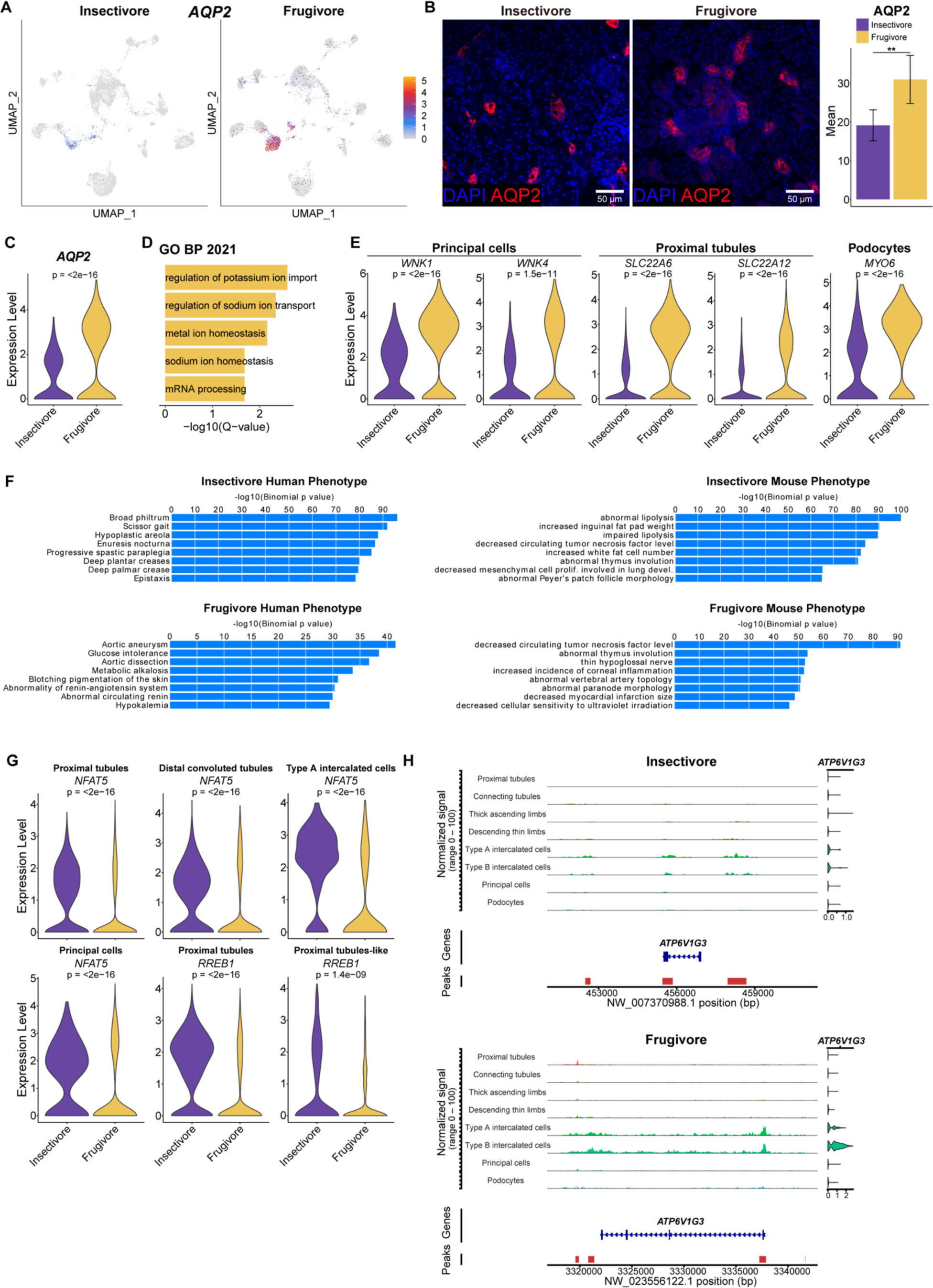
scRNA-seq and scATAC-seq analysis of bat kidney. (**A**) UMAPs of principal cell marker gene *AQP2* expression in each species. (**B**) (Left) Representative images of *SLC26A4* immunofluorescence (red) in bat kidneys. Nuclei are stained with DAPI (blue). (Right) Quantification of *AQP2* immunofluorescence normalized to nuclei in bat kidneys. Results represent arbitrary units of fluorescence (AU) mean ± standard error of the mean (SEM) derived from 3 insectivorous and 3 frugivorous bats (n = 3/phenotype, n = 10 images/individual [see Materials and Methods]). Mixed effects model \*\**p*-value = .001. (**C**) Violin plot of *AQP2* expression in principal cells. (**D**) Bar plots showing GO Biological Process 2021 terms enriched in frugivore principal cells. (**E**) Violin plots of differentially expressed genes in bat kidney cells. (**F**) Bar plots showing GREAT human and mouse phenotypes enriched in insectivore and frugivore kidneys. (**G**) Violin plots of differentially expressed TFs in bat renal epithelial cells. (**H**) scATAC-seq coverage plots of *ATP6V1G3* in bat kidneys. SCTransform-normalized expression plot visualized on the right by cell-type.

**Fig. S5.**
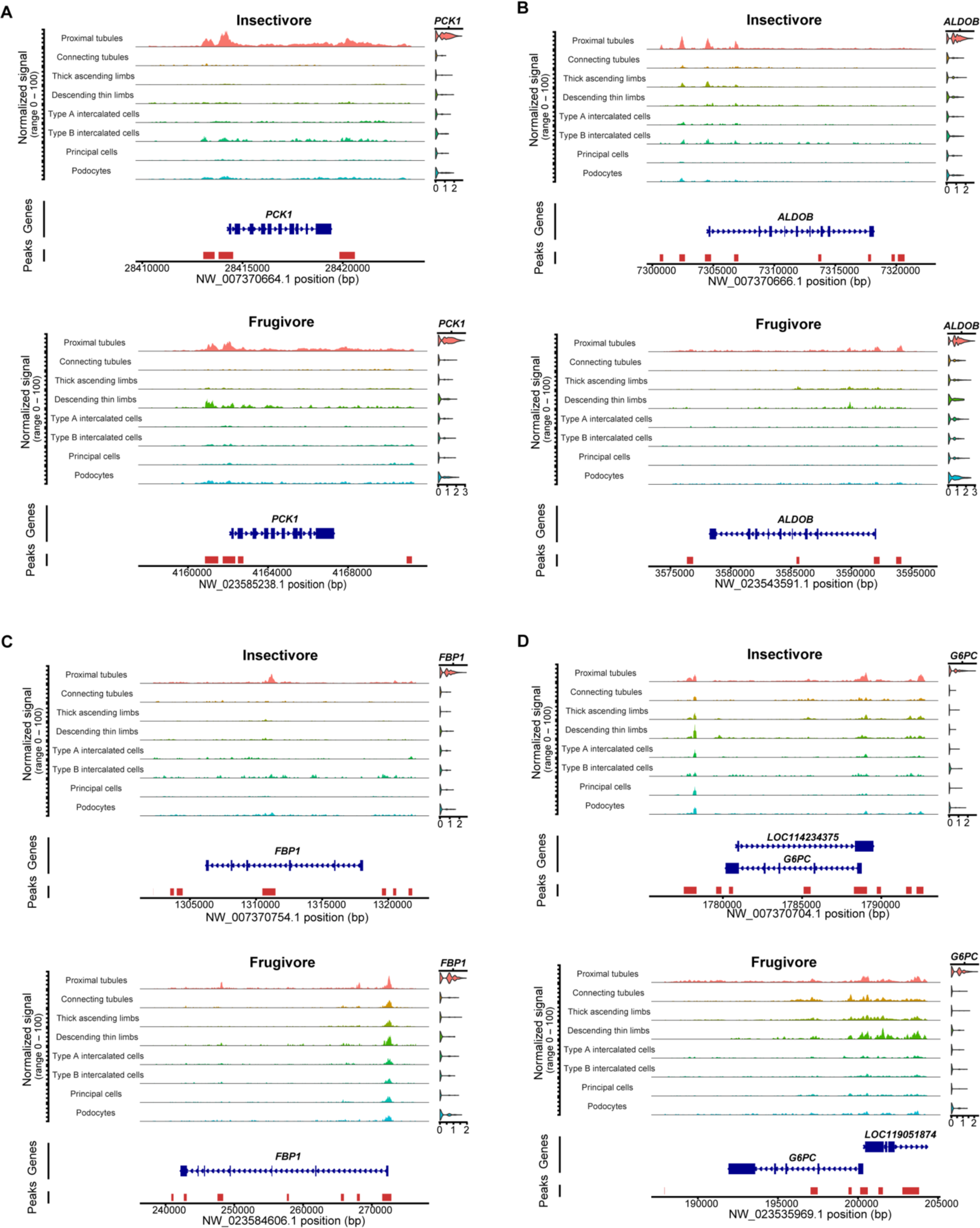
scATAC-seq coverage plots of diabetes-associated genes in bat kidneys. (**A-D**) scATAC-seq coverage plots of *PCK1* (**A**), *ALDOB* (**B**), *FBP1* (**C**) and *G6PC* (**D**) in bat kidneys. SCTransform-normalized expression plot visualized on the right by cell-type.

**Fig. S6.**
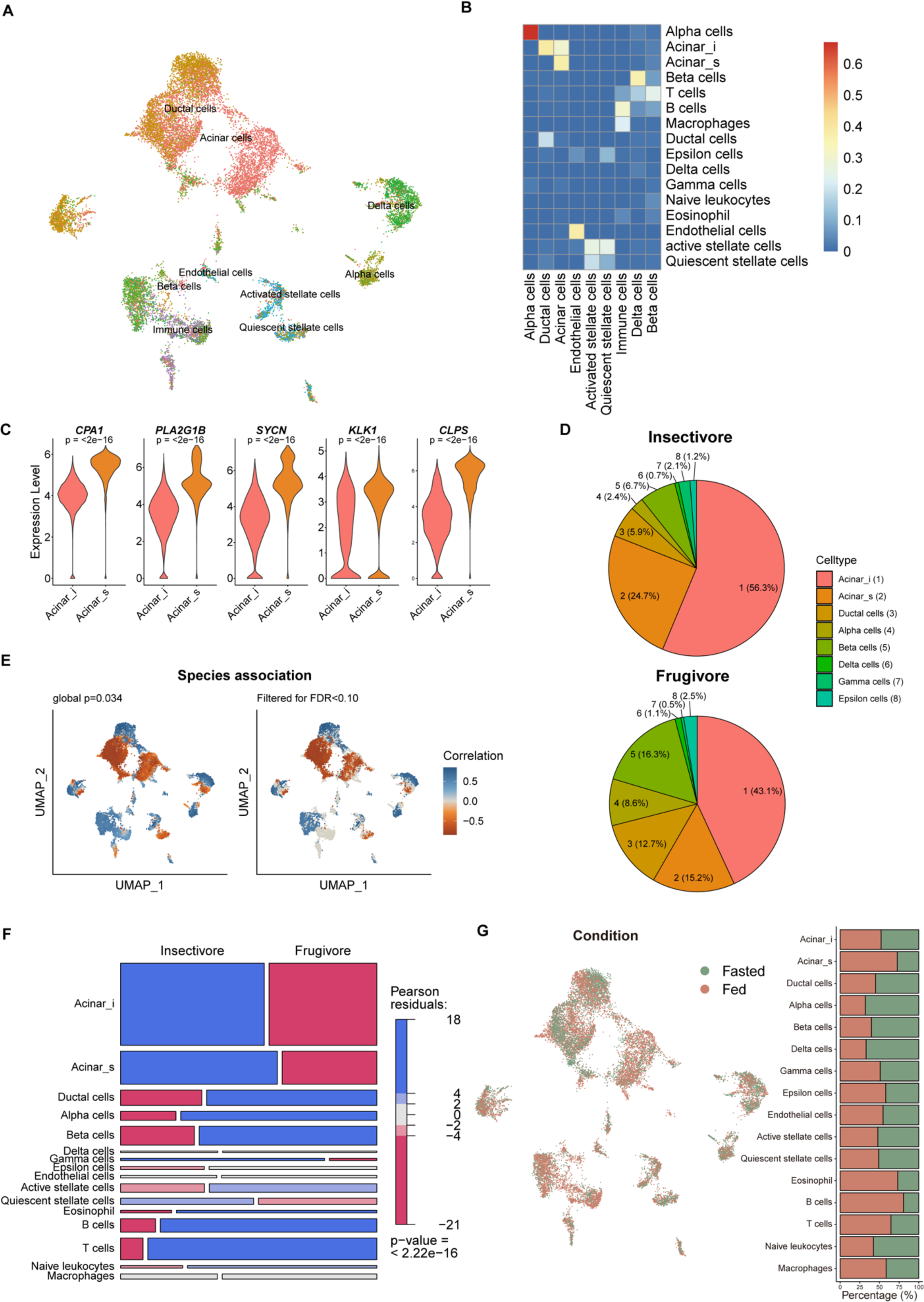
Single-cell composition analysis of bat pancreas. (**A**) UMAP of bat pancreas cell-types automatically annotated with human pancreas single-cell reference data from Azimuth (*42*). (**B**) Jaccard overlaps of auto-annotations in **A** (horizontal) with our integrated species annotations in Fig. 4B (vertical). (**C**) Violin plots of proximal tubule marker gene expression in bat proximal tubules and proximal tubules-like cells. (**D**) Pie charts of cell-type percentages across renal epithelial cells. (**E**) CNA of species associations to cell-types (**F**) Pearson residuals visualized by mosaic plot of all cell-types identified in bat pancreases. (**G**) UMAP of bat pancreases cell-types by condition. Bar chart shows the proportion per condition for each cell-type.

**Fig. S7.**
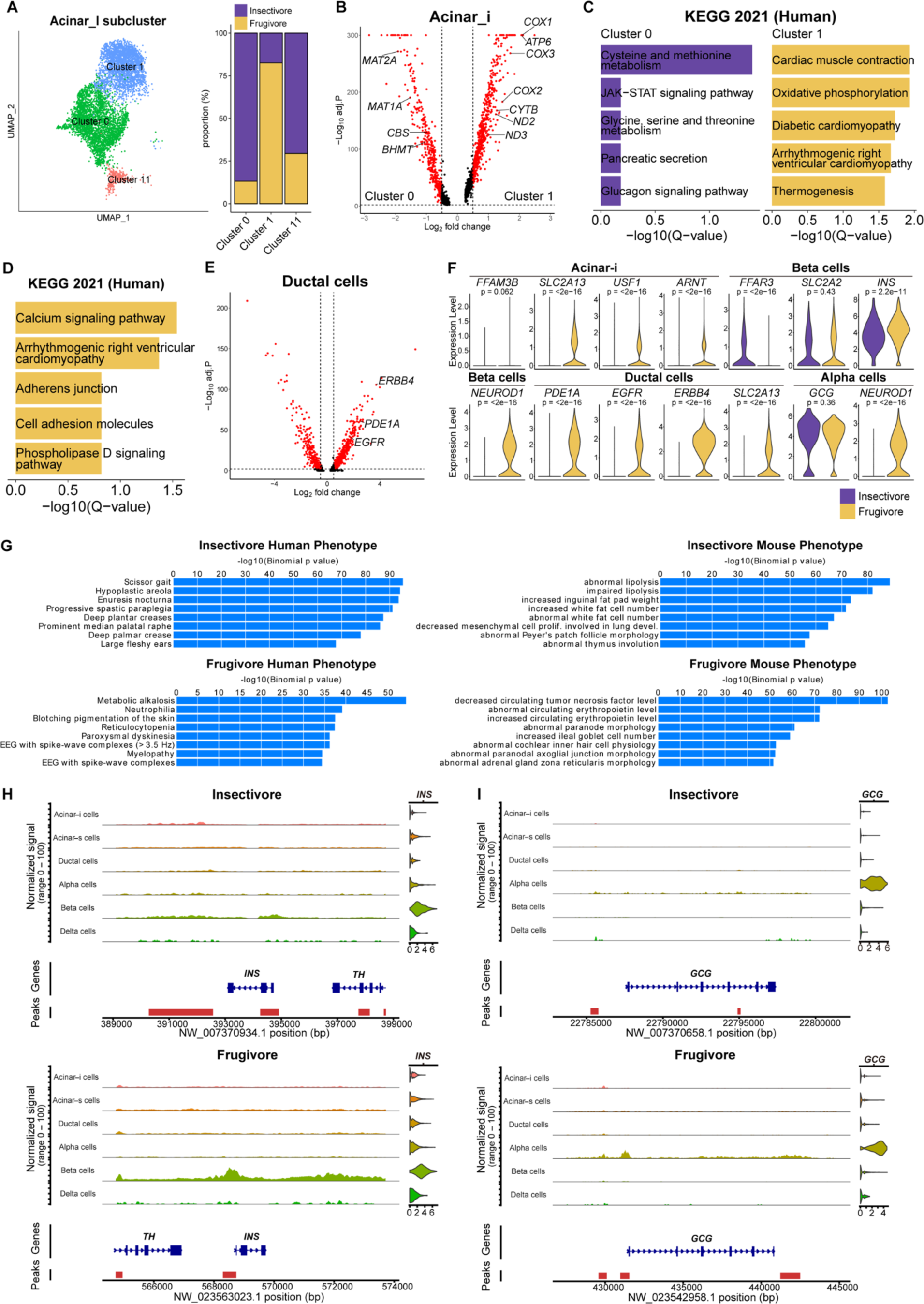
scRNA-seq and scATAC-seq analysis of bat pancreas. (**A**) (Left) UMAP of acinar-i subclusters. (Right) Proportion of species in each acinar-i subcluster. (**B**) Volcano plot showing differentially expressed genes between acinar-i subcluster 0 (insectivore-enriched) and subcluster 1 (frugivore-enriched). (**C**) Bar plots showing KEGG Human 2021 pathways enriched in acinar-i subcluster 0 (insectivore-enriched) and subcluster 1 (**D**) Bar plots showing KEGG Human 2021 pathways enriched in frugivore ductal cells. (**E**) Volcano plot showing differentially expressed genes between species in ductal cells. (**F**) Violin plots of differentially expressed genes, except *GCG*, in bat pancreas cells. (**G**) Bar plots showing GREAT human and mouse phenotypes enriched in insectivore and frugivore pancreases. (**H**) scATAC-seq coverage plots of *INS* in bat pancreases. SCTransform-normalized expression plot visualized on the right by cell-type. (**I**) scATAC-seq coverage plots of *GCG* in bat pancreases. SCTransform-normalized expression plot visualized on the right by cell-type.

**Fig. S8.**
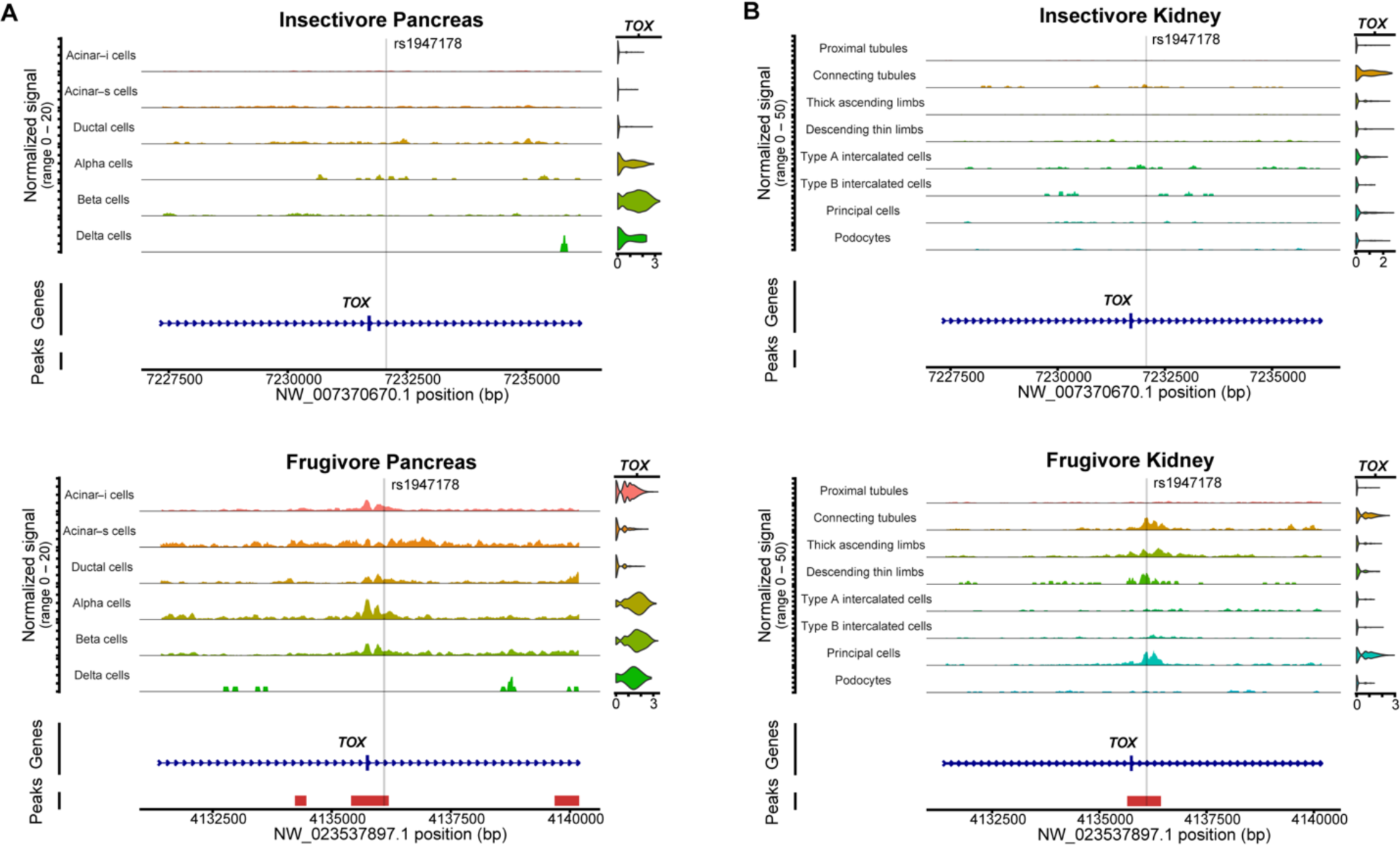
scATAC-seq coverage plots of diabetes-associated SNP rs1947178 in bats (151,857 bp and 171,812 bp downstream of *TOX* TSS in insectivorous and frugivorous bats, respectively). (**A-B**) scATAC-seq coverage plots of rs1947178 at *TOX* locus in bat pancreas (**A**) and kidney (**B**). SCTransform-normalized expression plot visualized on the right by cell-type.

**Table S1.** Sequencing QC. A1-2 = fasted fruit bats, A3-4 = treated fruit bats, K1-3 = treated insect bats, K4-5 = fasted insect bats, J1-2 = fasted fruit bats, J4= treated fruit bat, P1-2 = treated insect bats, P4-5 = fasted insect bats.

| **Sample** | **Total RNA read pairs** | **Mean RNA reads per cell** | **Total ATAC read pairs** | **Mean ATAC fragments per cell** |
| --- | --- | --- | --- | --- |
| A1 | 657490451 | 216921.9568 | 705761251 | 232847.6579 |
| A2 | 543890027 | 126251.1669 | 740382314 | 171862.1899 |
| A3 | 819522891 | 245734.0003 | 651414009 | 195326.5394 |
| A4 | 774977825 | 289278.7701 | 730688551 | 272746.7529 |
| E1 | 672912456 | 240411.7385 | 633695035 | 226400.5127 |
| E3 | 759040415 | 385887.3488 | 787887849 | 400553.0498 |
| E4 | 777840969 | 497022.9834 | 549534015 | 351139.9457 |
| E5 | 639507539 | 144717.7051 | 807937847 | 182832.733 |
| J1 | 642821902 | 305088.7053 | 726793113 | 344942.1514 |
| J2 | 559726040 | 200762.5681 | 719196923 | 257961.5936 |
| J4 | 578768902 | 123827.3218 | 949710083 | 203190.0049 |
| P1 | 927901678 | 548405.247 | 635548081 | 375619.4332 |
| P2 | 787957604 | 294014.0313 | 613644645 | 228971.8825 |
| P4 | 770059917 | 637994.9602 | 502313147 | 416166.6504 |
| P5 | 866020327 | 541601.2051 | 523034574 | 327101.0469 |
| **Mean** | 718562596.2 | 319861.3139 | 685169429.1 | 279177.4763 |
| **Std Dev.** | 114842825.9 | 165476.5497 | 117573079 | 82871.35383 |

**Table S2. Differentially expressed genes between fasted and fed states in kidney for each species.**

**Table S3. Differentially expressed genes between fasted and fed states in pancreas for each species.**

**Table S4.**
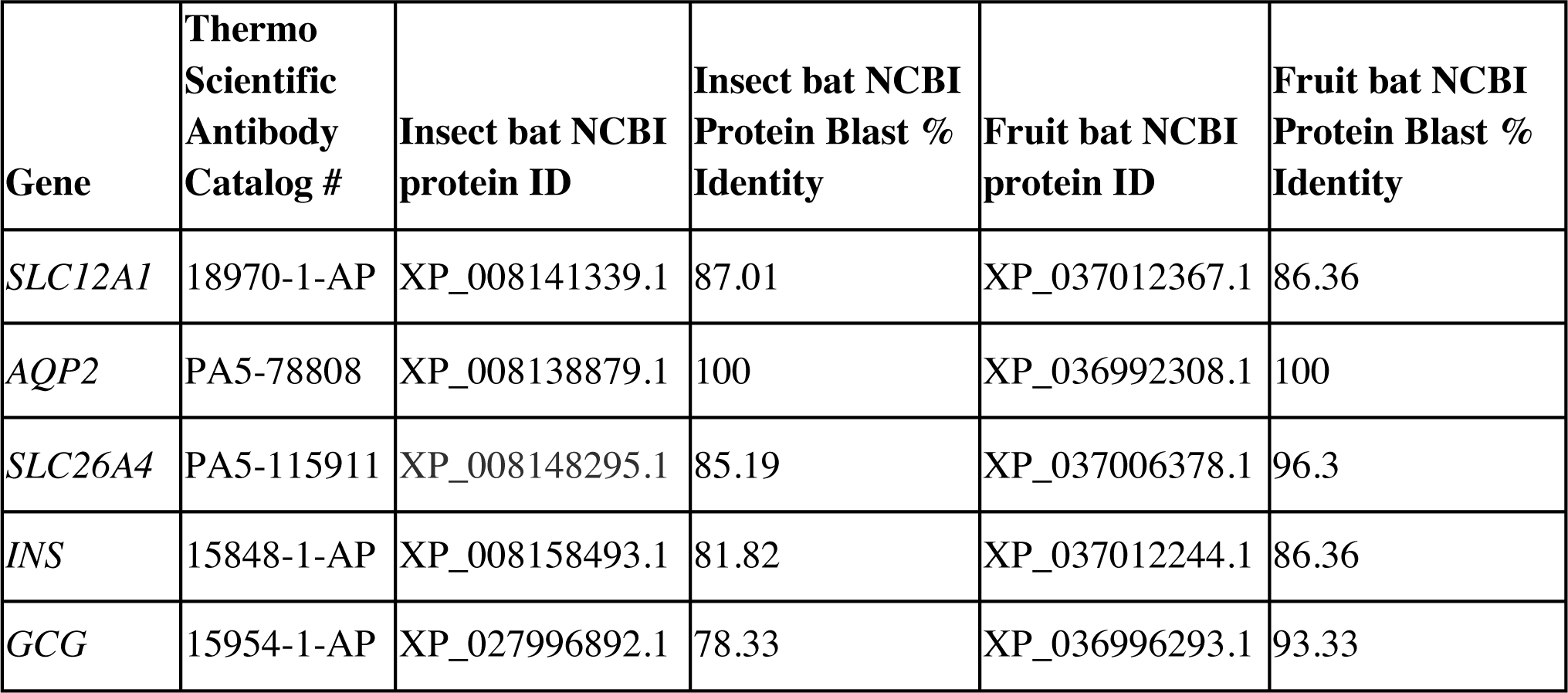
Immunofluorescence antibodies and epitope matching to bats.

**Table S5. Differentially expressed genes between species in the kidney.** Positive fold change indicates upregulation in frugivores.

**Table S6. Gene enrichment analyses of differentially expressed genes between species in the kidney.**

**Table S7. Cell-type-specific ATAC peaks in the insectivore kidney.**

**Table S8. Cell-type-specific ATAC peaks in the frugivore kidney.**

**Table S9. T1 and T2 diabetes-associated SNP list in *hg38* coordinates.**

**Table S10. T1 and T2 diabetes-associated SNP overlaps with bat kidney scATAC-seq peaks in *hg38* coordinates.**

**Table S11. Differentially expressed genes between species in the pancreas.** Positive fold change indicates upregulation in frugivores.

**Table S12. Gene enrichment analyses of differentially expressed genes between species in the pancreas.**

**Table S13. Cell-type-specific ATAC peaks in insectivore pancreas.**

**Table S14. Cell-type-specific ATAC peaks in frugivore pancreas.**

**Table S15. T1 and T2 diabetes-associated SNP overlaps with bat pancreas scATAC-seq peaks in *hg38* coordinates.**

**Table S16. Orthogroup IDs from Orthofinder for re-naming bat genome features.**

## References and Notes

1. T. H. Fleming, W. John Kress, A brief history of fruits and frugivores. Acta Oecol. 37, 521– 530 (2011).

2. G. F. Gunnell, N. B. Simmons, Fossil Evidence and the Origin of Bats. J. Mamm. Evol. 12, 209–246 (2005).

3. F. Meng, L. Zhu, W. Huang, D. M. Irwin, S. Zhang, Bats: Body mass index, forearm mass index, blood glucose levels and SLC2A2 genes for diabetes. Sci. Rep. 6, 29960 (2016).

4. A. Sadier, K. T. Davies, L. R. Yohe, K. Yun, P. Donat, B. P. Hedrick, E. R. Dumont, L. M. Dávalos, S. J. Rossiter, K. E. Sears, Multifactorial processes underlie parallel opsin loss in neotropical bats. Elife. 7 (2018), doi:10.7554/eLife.37412.

5. K. T. J. Davies, L. R. Yohe, J. Almonte, M. K. R. Sánchez, E. M. Rengifo, E. R. Dumont, K. E. Sears, L. M. Dávalos, S. J. Rossiter, Foraging shifts and visual preadaptation in ecologically diverse bats. Mol. Ecol. 29, 1839–1859 (2020).

6. K. Kshitish Acharya, A. Roy, A. Krishna, Relative role of olfactory cues and certain non-olfactory factors in foraging of fruit-eating bats. Behav. Processes. 44, 59–64 (1998).

7. F. Sánchez, C. Korine, M. Steeghs, L.-J. Laarhoven, S. M. Cristescu, F. J. M. Harren, R. Dudley, B. Pinshow, Ethanol and methanol as possible odor cues for Egyptian fruit bats (Rousettus aegyptiacus). J. Chem. Ecol. 32, 1289–1300 (2006).

8. E. R. Dumont, Cranial shape in fruit, nectar, and exudate feeders: implications for interpreting the fossil record. Am. J. Phys. Anthropol. 102, 187–202 (1997).

9. M. R. Marchán-Rivadeneira, C. J. Phillips, R. E. Strauss, J. Antonio Guerrero, C. A. Mancina, R. J. Baker, Cranial differentiation of fruit-eating bats (GenusArtibeus) based on size-standardized data. Acta Chiropt. 12, 143–154 (2010).

10. E. R. Dumont, K. Samadevam, I. Grosse, O. M. Warsi, B. Baird, L. M. Davalos, Selection for mechanical advantage underlies multiple cranial optima in new world leaf-nosed bats. Evolution. 68, 1436–1449 (2014).

11. J. H. Arbour, A. A. Curtis, S. E. Santana, Signatures of echolocation and dietary ecology in the adaptive evolution of skull shape in bats. Nat. Commun. 10, 1–13 (2019).

12. L. V. García-Herrera, L. A. Ramírez-Fráncel, G. Guevara, G. Reinoso-Flórez, A. Sánchez-Hernández, B. K. Lim, S. Losada-Prado, Foraging strategies, craniodental traits, and interaction in the bite force of Neotropical frugivorous bats (Phyllostomidae: Stenodermatinae). Ecol. Evol. 11, 13756–13772 (2021).

13. D. Massoud, M. M. A. Abumandour, Anatomical features of the tongue of two chiropterans endemic in the Egyptian fauna; the Egyptian fruit bat (Rousettus aegyptiacus) and insectivorous bat (Pipistrellus kuhlii). Acta Histochem. 122, 151503 (2020).

14. A. N. Makanya, J. N. Maina, T. M. Mayhew, S. A. Tschanz, P. H. Burri, A stereological comparison of villous and microvillous surfaces in small intestines of frugivorous and entomophagous bats: species, inter-individual and craniocaudal differences. J. Exp. Biol. 200, 2415–2423 (1997).

15. R. Gadelha-Alves, A. M. D. S. Rozensztranch, O. Rocha-Barbosa, Comparative Intestinal Histomorphology of Five Species of Phyllostomid Bats (Phyllostomidae, Microchiroptera): Ecomorphological Relations with Alimentary Habits. Int. J. Morphol. 26, 591–602 (2008).

16. J. E. Schondube, L. G. Herrera-M, C. Martínez del Rio, Diet and the evolution of digestion and renal function in phyllostomid bats. Zoology . 104, 59–73 (2001).

17. G. Casotti, L. Gerardo Herrera M, J. J. Flores M, C. A. Mancina, E. J. Braun, Relationships between renal morphology and diet in 26 species of new world bats (suborder microchiroptera). Zoology . 109, 196–207 (2006).

18. A. J. Michelmore, D. J. Keegan, B. Kramer, Immunocytochemical Identification of Endocrine Cells in the Pancreas of the Fruit Bat,Rousettus aegyptiacus. General and Comparative Endocrinology. 110 (1998), pp. 319–325.

19. A. O. P. Protzek, A. Rafacho, B. A. Viscelli, J. R. Bosqueiro, A. P. Cappelli, F. M. M. Paula, A. C. Boschero, E. C. Pinheiro, Insulin and glucose sensitivity, insulin secretion and β-cell distribution in endocrine pancreas of the fruit bat Artibeus lituratus. Comp. Biochem. Physiol. A Mol. Integr. Physiol. 157, 142–148 (2010).

20. V. Sharma, N. Hecker, J. G. Roscito, L. Foerster, B. E. Langer, M. Hiller, A genomics approach reveals insights into the importance of gene losses for mammalian adaptations. Nat. Commun. 9, 1215 (2018).

21. K. Wang, S. Tian, J. Galindo-González, L. M. Dávalos, Y. Zhang, H. Zhao, Molecular adaptation and convergent evolution of frugivory in Old World and neotropical fruit bats. Mol. Ecol. 29, 4366–4381 (2020).

22. D. J. Keegan, Aspects of the assimilation of sugars by Rousettus aegyptiacus. Comp. Biochem. Physiol. A Physiol. 58, 349–352 (1977).

23. M. B. Freitas, J. F. Queiroz, C. I. Dias Gomes, C. B. Collares-Buzato, H. C. Barbosa, A. C. Boschero, C. A. Gonçalves, E. C. Pinheiro, Reduced insulin secretion and glucose intolerance are involved in the fasting susceptibility of common vampire bats. Gen. Comp. Endocrinol. 183, 1–6 (2013).

24. X. Peng, X. He, Y. Sun, J. Liang, H. Xie, Difference in glucose tolerance between phytophagous and insectivorous bats. Journal of Comparative Physiology. B, Biochemical, Systemic, and Environmental Physiology; Heidelberg. 189, 751–756 (2019).

25. O. Amitai, S. Holtze, S. Barkan, E. Amichai, C. Korine, B. Pinshow, C. C. Voigt, Fruit bats (Pteropodidae) fuel their metabolism rapidly and directly with exogenous sugars. J. Exp. Biol. 213, 2693–2699 (2010).

26. J. E. Campbell, C. B. Newgard, Mechanisms controlling pancreatic islet cell function in insulin secretion. Nat. Rev. Mol. Cell Biol. 22, 142–158 (2021).

27. M. Karpińska, M. Czauderna, Pancreas-Its Functions, Disorders, and Physiological Impact on the Mammals’ Organism. Front. Physiol. 13, 807632 (2022).

28. M. S. Balzer, T. Rohacs, K. Susztak, How Many Cell Types Are in the Kidney and What Do They Do? Annu. Rev. Physiol. 84, 507–531 (2022).

29. I. H. de Boer, K. M. Utzschneider, The kidney’s role in systemic metabolism-still much to learn. Nephrol. Dial. Transplant. 32, 588–590 (2017).

30. Z. Arad, C. Korine, Effect of water restriction on energy and water balance and osmoregulation of the fruit bat Rousettus aegyptiacus. J. Comp. Physiol. B. 163, 401–405 (1993).

31. H. Xu, Y. Liu, F. Meng, B. He, N. Han, G. Li, S. J. Rossiter, S. Zhang, Multiple bursts of pancreatic ribonuclease gene duplication in insect-eating bats. Gene. 526, 112–117 (2013).

32. M. C. Janiak, M. E. Chaney, A. J. Tosi, Evolution of Acidic Mammalian Chitinase Genes (CHIA) Is Related to Body Mass and Insectivory in Primates. Mol. Biol. Evol. 35, 607–622 (2018).

33. Y. Liu, H. Xu, X. Yuan, S. J. Rossiter, S. Zhang, Multiple adaptive losses of alanine-glyoxylate aminotransferase mitochondrial targeting in fruit-eating bats. Mol. Biol. Evol. 29, 1507–1511 (2012).

34. B. Shen, X. Han, J. Zhang, S. J. Rossiter, S. Zhang, Adaptive evolution in the glucose transporter 4 gene Slc2a4 in Old World fruit bats (family: Pteropodidae). PLoS One. 7, e33197 (2012).

35. L. Fang, B. Shen, D. M. Irwin, S. Zhang, Parallel evolution of the glycogen synthase 1 (muscle) gene Gys1 between Old World and New World fruit bats (Order: Chiroptera). Biochem. Genet. 52, 443–458 (2014).

36. Y. Qian, T. Fang, B. Shen, S. Zhang, The glycogen synthase 2 gene (Gys2) displays parallel evolution between Old World and New World fruit bats. J. Mol. Evol. 78, 66–74 (2014).

37. B. Shen, T. Fang, T. Yang, G. Jones, D. M. Irwin, S. Zhang, Relaxed evolution in the tyrosine aminotransferase gene tat in old world fruit bats (Chiroptera: Pteropodidae). PLoS One. 9, e97483 (2014).

38. L. Zhu, Q. Yin, D. M. Irwin, S. Zhang, Phosphoenolpyruvate carboxykinase 1 gene (Pck1) displays parallel evolution between Old World and New World fruit bats. PLoS One. 10, e0118666 (2015).

39. Q. Yin, L. Zhu, D. Liu, D. M. Irwin, S. Zhang, Y.-H. Pan, Molecular Evolution of the Nuclear Factor (Erythroid-Derived 2)-Like 2 Gene Nrf2 in Old World Fruit Bats (Chiroptera: Pteropodidae). PLoS One. 11, e0146274 (2016).

40. I. Agnarsson, C. M. Zambrana-Torrelio, N. P. Flores-Saldana, L. J. May-Collado, A time-calibrated species-level phylogeny of bats (Chiroptera, Mammalia). PLoS Curr. 3, RRN1212 (2011).

41. M. Laska, Food transit times and carbohydrate use in three phyllostomid bat species. Zeitschrift für Säugetierkunde. 55, 49–54 (1990).

42. Y. Hao, S. Hao, E. Andersen-Nissen, W. M. Mauck 3rd, S. Zheng, A. Butler, M. J. Lee, A. J. Wilk, C. Darby, M. Zager, P. Hoffman, M. Stoeckius, E. Papalexi, E. P. Mimitou, J. Jain, A. Srivastava, T. Stuart, L. M. Fleming, B. Yeung, A. J. Rogers, J. M. McElrath, C. A. Blish, R. Gottardo, P. Smibert, R. Satija, Integrated analysis of multimodal single-cell data. Cell. 184, 3573–3587.e29 (2021).

43. L. Montefiori, L. Hernandez, Z. Zhang, Y. Gilad, C. Ober, G. Crawford, M. Nobrega, N. Jo Sakabe, Reducing mitochondrial reads in ATAC-seq using CRISPR/Cas9. Sci. Rep. 7, 2451 (2017).

44. J.-J. Jin, W.-B. Yu, J.-B. Yang, Y. Song, C. W. dePamphilis, T.-S. Yi, D.-Z. Li, GetOrganelle: a fast and versatile toolkit for accurate de novo assembly of organelle genomes. Genome Biol. 21, 241 (2020).

45. D. M. Emms, S. Kelly, OrthoFinder: phylogenetic orthology inference for comparative genomics. Genome Biol. 20, 238 (2019).

46. T. Stuart, A. Srivastava, S. Madad, C. A. Lareau, R. Satija, Single-cell chromatin state analysis with Signac. Nat. Methods. 18, 1333–1341 (2021).

47. I. Korsunsky, N. Millard, J. Fan, K. Slowikowski, F. Zhang, K. Wei, Y. Baglaenko, M. Brenner, P.-R. Loh, S. Raychaudhuri, Fast, sensitive and accurate integration of single-cell data with Harmony. Nat. Methods. 16, 1289–1296 (2019).

48. S. Álvarez-Carretero, A. U. Tamuri, M. Battini, F. F. Nascimento, E. Carlisle, R. J. Asher, Z. Yang, P. C. J. Donoghue, M. Dos Reis, A species-level timeline of mammal evolution integrating phylogenomic data. Nature. 602, 263–267 (2022).

49. R. A. Haeusler, K. Hartil, B. Vaitheesvaran, I. Arrieta-Cruz, C. M. Knight, J. R. Cook, H. L. Kammoun, M. A. Febbraio, R. Gutierrez-Juarez, I. J. Kurland, D. Accili, Integrated control of hepatic lipogenesis versus glucose production requires FoxO transcription factors. Nat. Commun. 5, 5190 (2014).

50. C. Q. Hoang, M. A. Hale, A. C. Azevedo-Pouly, H. P. Elsässer, T. G. Deering, S. G. Willet, F. C. Pan, M. A. Magnuson, C. V. E. Wright, G. H. Swift, R. J. MacDonald, Transcriptional Maintenance of Pancreatic Acinar Identity, Differentiation, and Homeostasis by PTF1A. Mol. Cell. Biol. 36, 3033–3047 (2016).

51. M. A. Fenech, C. M. Sullivan, L. T. Ferreira, R. Mehmood, W. A. MacDonald, P. B. Stathopulos, C. L. Pin, Atp2c2 Is Transcribed From a Unique Transcriptional Start Site in Mouse Pancreatic Acinar Cells. J. Cell. Physiol. 231, 2768–2778 (2016).

52. Y. A. Reshef, L. Rumker, J. B. Kang, A. Nathan, I. Korsunsky, S. Asgari, M. B. Murray, D. B. Moody, S. Raychaudhuri, Co-varying neighborhood analysis identifies cell populations associated with phenotypes of interest from single-cell transcriptomics. Nat. Biotechnol. 40, 355–363 (2022).

53. O. Franzén, L.-M. Gan, J. L. M. Björkegren, PanglaoDB: a web server for exploration of mouse and human single-cell RNA sequencing data. Database . 2019 (2019), doi:10.1093/database/baz046.

54. Z. Miao, M. S. Balzer, Z. Ma, H. Liu, J. Wu, R. Shrestha, T. Aranyi, A. Kwan, A. Kondo, M. Pontoglio, J. Kim, M. Li, K. H. Kaestner, K. Susztak, Single cell regulatory landscape of the mouse kidney highlights cellular differentiation programs and disease targets. Nat. Commun. 12, 2277 (2021).

55. Y. Muto, P. C. Wilson, N. Ledru, H. Wu, H. Dimke, S. S. Waikar, B. D. Humphreys, Single cell transcriptional and chromatin accessibility profiling redefine cellular heterogeneity in the adult human kidney. Nat. Commun. 12, 2190 (2021).

56. B. B. Lake, R. Menon, S. Winfree, Q. Hu, R. M. Ferreira, K. Kalhor, D. Barwinska, E. A. Otto, M. Ferkowicz, D. Diep, N. Plongthongkum, A. Knoten, S. Urata, A. S. Naik, S. Eddy, B. Zhang, Y. Wu, D. Salamon, J. C. Williams, X. Wang, K. S. Balderrama, P. Hoover, E. Murray, A. Vijayan, F. Chen, S. S. Waikar, S. Rosas, F. P. Wilson, P. M. Palevsky, K. Kiryluk, J. R. Sedor, R. D. Toto, C. Parikh, E. H. Kim, E. Z. Macosko, P. V. Kharchenko, J. P. Gaut, J. B. Hodgin, M. T. Eadon, P. C. Dagher, T. M. El-Achkar, K. Zhang, M. Kretzler, S. Jain, for the KPMP consortium, An atlas of healthy and injured cell states and niches in the human kidney. bioRxiv (2021), p. 2021.07.28.454201.

57. L. Chen, J. Z. Clark, J. W. Nelson, B. Kaissling, D. H. Ellison, M. A. Knepper, Renal-Tubule Epithelial Cell Nomenclature for Single-Cell RNA-Sequencing Studies. J. Am. Soc. Nephrol. 30, 1358–1364 (2019).

58. E. P. Paksuz, Renal adaptation in relation to insectivorous feeding habit in the greater mouse-eared bat, Myotis myotis (Chiroptera: Vespertilionidae). Anat. Rec. (2022), doi:10.1002/ar.24946.

59. A. Roy, M. M. Al-bataineh, N. M. Pastor-Soler, Collecting duct intercalated cell function and regulation. Clin. J. Am. Soc. Nephrol. 10, 305–324 (2015).

60. S. A. Lanham-New, The balance of bone health: tipping the scales in favor of potassium-rich, bicarbonate-rich foods. J. Nutr. 138, 172S–177S (2008).

61. A. Staruschenko, Regulation of transport in the connecting tubule and cortical collecting duct. Compr. Physiol. 2, 1541–1584 (2012).

62. P. C. Wilson, H. Wu, Y. Kirita, K. Uchimura, N. Ledru, H. G. Rennke, P. A. Welling, S. S. Waikar, B. D. Humphreys, The single-cell transcriptomic landscape of early human diabetic nephropathy. Proc. Natl. Acad. Sci. U. S. A. 116, 19619–19625 (2019).

63. N. Uehara-Watanabe, N. Okuno-Ozeki, A. Minamida, I. Nakamura, T. Nakata, K. Nakai, A. Yagi-Tomita, T. Ida, K. Ikeda, T. Kitani, N. Yamashita, M. Kamezaki, Y. Kirita, S. Matoba, K. Tamagaki, T. Kusaba, Direct evidence of proximal tubular proliferation in early diabetic nephropathy. Sci. Rep. 12, 778 (2022).

64. T. Furusho, S. Uchida, E. Sohara, The WNK signaling pathway and salt-sensitive hypertension. Hypertens. Res. 43, 733–743 (2020).

65. A. R. Rodan, A. Jenny, WNK Kinases in Development and Disease. Curr. Top. Dev. Biol. 123, 1–47 (2017).

66. W. H. Yiu, D. W. L. Wong, L. Y. Y. Chan, J. C. K. Leung, K. W. Chan, H. Y. Lan, K. N. Lai, S. C. W. Tang, Tissue kallikrein mediates pro-inflammatory pathways and activation of protease-activated receptor-4 in proximal tubular epithelial cells. PLoS One. 9, e88894 (2014).

67. P. Madeddu, C. Emanueli, S. El-Dahr, Mechanisms of disease: the tissue kallikrein-kinin system in hypertension and vascular remodeling. Nat. Clin. Pract. Nephrol. 3, 208–221 (2007).

68. G. Colla, H.-J. Kim, M. C. Kyriacou, Y. Rouphael, Nitrate in fruits and vegetables. Sci. Hortic. . 237, 221–238 (2018).

69. S. El Moghrabi, P. Houillier, N. Picard, F. Sohet, B. Wootla, M. Bloch-Faure, F. Leviel, L. Cheval, S. Frische, P. Meneton, D. Eladari, R. Chambrey, Tissue kallikrein permits early renal adaptation to potassium load. Proc. Natl. Acad. Sci. U. S. A. 107, 13526–13531 (2010).

70. D. Szklarczyk, A. Franceschini, S. Wyder, K. Forslund, D. Heller, J. Huerta-Cepas, M. Simonovic, A. Roth, A. Santos, K. P. Tsafou, M. Kuhn, P. Bork, L. J. Jensen, C. von Mering, STRING v10: protein-protein interaction networks, integrated over the tree of life. Nucleic Acids Res. 43, D447–52 (2015).

71. D. B. Mount, Thick ascending limb of the loop of Henle. Clin. J. Am. Soc. Nephrol. 9, 1974–1986 (2014).

72. G. Schiano, B. Glaudemans, E. Olinger, N. Goelz, M. Müller, D. Loffing-Cueni, G. Deschenes, J. Loffing, O. Devuyst, The Urinary Excretion of Uromodulin is Regulated by the Potassium Channel ROMK. Sci. Rep. 9, 19517 (2019).

73. E. H. Studier, D. E. Wilson, Natural urine concentrations and composition in neotropical bats. Comp. Biochem. Physiol. A Physiol. 75, 509–515 (1983).

74. J. N. Lorenz, N. R. Baird, L. M. Judd, W. T. Noonan, A. Andringa, T. Doetschman, P. A. Manning, L. H. Liu, M. L. Miller, G. E. Shull, Impaired Renal NaCl Absorption in Mice Lacking the ROMK Potassium Channel, a Model for Type II Bartter’s Syndrome*. J. Biol. Chem. 277, 37871–37880 (2002).

75. S.-H. Lin, I.-S. Yu, S.-T. Jiang, S.-W. Lin, P. Chu, A. Chen, H.-K. Sytwu, E. Sohara, S. Uchida, S. Sasaki, S.-S. Yang, Impaired phosphorylation of Na(+)-K(+)-2Cl(-) cotransporter by oxidative stress-responsive kinase-1 deficiency manifests hypotension and Bartter-like syndrome. Proc. Natl. Acad. Sci. U. S. A. 108, 17538–17543 (2011).

76. J. Xue, L. Thomas, J. D. Rieg, T. Rieg, Generation and characterization of thick ascending limb-specific NHE3 knockout mice. FASEB J. 34, 1–1 (2020).

77. D. B. Simon, F. E. Karet, J. M. Hamdan, A. DiPietro, S. A. Sanjad, R. P. Lifton, Bartter’s syndrome, hypokalaemic alkalosis with hypercalciuria, is caused by mutations in the Na-K-2Cl cotransporter NKCC2. Nat. Genet. 13, 183–188 (1996).

78. D. B. Simon, F. E. Karet, J. Rodriguez-Soriano, J. H. Hamdan, A. DiPietro, H. Trachtman, S. A. Sanjad, R. P. Lifton, Genetic heterogeneity of Bartter’s syndrome revealed by mutations in the K+ channel, ROMK. Nat. Genet. 14, 152–156 (1996).

79. N. Jeck, C. Derst, E. Wischmeyer, H. Ott, S. Weber, C. Rudin, H. W. Seyberth, J. Daut, A. Karschin, M. Konrad, Functional heterogeneity of ROMK mutations linked to hyperprostaglandin E syndrome. Kidney Int. 59, 1803–1811 (2001).

80. B. Shen, X. Han, G. Jones, S. J. Rossiter, S. Zhang, Adaptive evolution of the myo6 gene in old world fruit bats (family: pteropodidae). PLoS One. 8, e62307 (2013).

81. N. Otani, M. Ouchi, K. Hayashi, P. Jutabha, N. Anzai, Roles of organic anion transporters (OATs) in renal proximal tubules and their localization. Anat. Sci. Int. 92, 200–206 (2017).

82. R. T. Keenan, The biology of urate. Semin. Arthritis Rheum. 50, S2–S10 (2020).

83. B. Jakše, B. Jakše, M. Pajek, J. Pajek, Uric Acid and Plant-Based Nutrition. Nutrients. 11 (2019), doi:10.3390/nu11081736.

84. Y. Chino, Y. Samukawa, S. Sakai, Y. Nakai, J.-I. Yamaguchi, T. Nakanishi, I. Tamai, SGLT2 inhibitor lowers serum uric acid through alteration of uric acid transport activity in renal tubule by increased glycosuria. Biopharm. Drug Dispos. 35, 391–404 (2014).

85. I. D. Weiner, Roles of renal ammonia metabolism other than in acid-base homeostasis. Pediatr. Nephrol. 32, 933–942 (2017).

86. N. Gotoh, Q. Yan, Z. Du, D. Biemesderfer, M. Kashgarian, M. S. Mooseker, T. Wang, Altered renal proximal tubular endocytosis and histology in mice lacking myosin-VI. Cytoskeleton . 67, 178–192 (2010).

87. P. C. Wilson, Y. Muto, H. Wu, A. Karihaloo, S. S. Waikar, B. D. Humphreys, Multimodal single cell sequencing implicates chromatin accessibility and genetic background in diabetic kidney disease progression. Nat. Commun. 13, 5253 (2022).

88. V. Vallon, Molecular determinants of renal glucose reabsorption. Focus on “Glucose transport by human renal Na+/D-glucose cotransporters SGLT1 and SGLT2.” Am. J. Physiol. Cell Physiol. 300 (2011), pp. C6–8.

89. Y. Zhang, T. Liu, C. A. Meyer, J. Eeckhoute, D. S. Johnson, B. E. Bernstein, C. Nusbaum, R. M. Myers, M. Brown, W. Li, X. S. Liu, Model-based analysis of ChIP-Seq (MACS). Genome Biol. 9, R137 (2008).

90. V. A. Schneider, T. Graves-Lindsay, K. Howe, N. Bouk, H.-C. Chen, P. A. Kitts, T. D. Murphy, K. D. Pruitt, F. Thibaud-Nissen, D. Albracht, R. S. Fulton, M. Kremitzki, V. Magrini, C. Markovic, S. McGrath, K. M. Steinberg, K. Auger, W. Chow, J. Collins, G. Harden, T. Hubbard, S. Pelan, J. T. Simpson, G. Threadgold, J. Torrance, J. M. Wood, L. Clarke, S. Koren, M. Boitano, P. Peluso, H. Li, C.-S. Chin, A. M. Phillippy, R. Durbin, R. K. Wilson, P. Flicek, E. E. Eichler, D. M. Church, Evaluation of GRCh38 and de novo haploid genome assemblies demonstrates the enduring quality of the reference assembly. Genome Res. 27, 849–864 (2017).

91. C. Y. McLean, D. Bristor, M. Hiller, S. L. Clarke, B. T. Schaar, C. B. Lowe, A. M. Wenger, G. Bejerano, GREAT improves functional interpretation of cis-regulatory regions. Nat. Biotechnol. 28, 495–501 (2010).

92. P. Machanick, T. L. Bailey, MEME-ChIP: motif analysis of large DNA datasets. Bioinformatics. 27, 1696–1697 (2011).

93. Y. Wang, G. Jarad, P. Tripathi, M. Pan, J. Cunningham, D. R. Martin, H. Liapis, J. H. Miner, F. Chen, Activation of NFAT signaling in podocytes causes glomerulosclerosis. J. Am. Soc. Nephrol. 21, 1657–1666 (2010).

94. Y. Maeoka, Y. Wu, T. Okamoto, S. Kanemoto, X. P. Guo, A. Saito, R. Asada, K. Matsuhisa, T. Masaki, K. Imaizumi, M. Kaneko, NFAT5 up-regulates expression of the kidney-specific ubiquitin ligase gene Rnf183 under hypertonic conditions in inner-medullary collecting duct cells. J. Biol. Chem. 294, 101–115 (2019).

95. C. E. Pierreux, J. Stafford, D. Demonte, D. K. Scott, J. Vandenhaute, R. M. O’Brien, D. K. Granner, G. G. Rousseau, F. P. Lemaigre, Antiglucocorticoid activity of hepatocyte nuclear factor-6. Proc. Natl. Acad. Sci. U. S. A. 96, 8961–8966 (1999).

96. K. Yamamoto, T.-A. Matsuoka, S. Kawashima, S. Takebe, F. Kubo, T. Miyatsuka, H. Kaneto, I. Shimomura, A novel function of Onecut1 protein as a negative regulator of MafA gene expression. J. Biol. Chem. 288, 21648–21658 (2013).

97. A. Philippi, S. Heller, I. G. Costa, V. Senée, M. Breunig, Z. Li, G. Kwon, R. Russell, A. Illing, Q. Lin, M. Hohwieler, A. Degavre, P. Zalloua, S. Liebau, M. Schuster, J. Krumm, X. Zhang, R. Geusz, J. R. Benthuysen, A. Wang, J. Chiou, K. Gaulton, H. Neubauer, E. Simon, T. Klein, M. Wagner, G. Nair, C. Besse, C. Dandine-Roulland, R. Olaso, J.-F. Deleuze, B. Kuster, M. Hebrok, T. Seufferlein, M. Sander, B. O. Boehm, F. Oswald, M. Nicolino, C. Julier, A. Kleger, Mutations and variants of ONECUT1 in diabetes. Nat. Med. 27, 1928– 1940 (2021).

98. S. Prudente, F. Andreozzi, L. Mercuri, F. Alberico, A. Di Giamberardino, G. C. Mannino, O. Ludovico, P. Piscitelli, R. Di Paola, S. Morano, G. Penno, M. Carella, S. De Cosmo, V. Trischitta, F. Barbetti, Contribution of ONECUT1 variants to different forms of non-autoimmune diabetes mellitus in Italian patients. Acta Diabetol. 59, 1113–1116 (2022).

99. K. Ueda, K. Fujiki, K. Shirahige, C. E. Gomez-Sanchez, T. Fujita, M. Nangaku, M. Nagase, Genome-wide analysis of murine renal distal convoluted tubular cells for the target genes of mineralocorticoid receptor. Biochem. Biophys. Res. Commun. 445, 132–137 (2014).

100. E. Gomez-Sanchez, C. E. Gomez-Sanchez, The multifaceted mineralocorticoid receptor. Compr. Physiol. 4, 965–994 (2014).

101. M. Epstein, C. P. Kovesdy, C. M. Clase, M. M. Sood, R. Pecoits-Filho, Aldosterone, Mineralocorticoid Receptor Activation, and CKD: A Review of Evolving Treatment Paradigms. Am. J. Kidney Dis. 80, 658–666 (2022).

102. H. Joshi, B. Vastrad, N. Joshi, C. Vastrad, Integrated bioinformatics analysis reveals novel key biomarkers in diabetic nephropathy. SAGE Open Med. 10, 20503121221137005 (2022).

103. J. A. Bonomo, M. Guan, M. C. Y. Ng, N. D. Palmer, P. J. Hicks, J. M. Keaton, J. P. Lea, C. D. Langefeld, B. I. Freedman, D. W. Bowden, The ras responsive transcription factor RREB1 is a novel candidate gene for type 2 diabetes associated end-stage kidney disease. Hum. Mol. Genet. 23, 6441–6447 (2014).

104. F. Rothenberger, A. Velic, P. A. Stehberger, J. Kovacikova, C. A. Wagner, Angiotensin II stimulates vacuolar H+ -ATPase activity in renal acid-secretory intercalated cells from the outer medullary collecting duct. J. Am. Soc. Nephrol. 18, 2085–2093 (2007).

105. Y. Wang, T. He, J. G. Herman, E. Linghu, Y. Yang, F. Fuks, F. Zhou, L. Song, M. Guo, Methylation of ZNF331 is an independent prognostic marker of colorectal cancer and promotes colorectal cancer growth. Clin. Epigenetics. 9, 115 (2017).

106. A. Buniello, J. A. L. MacArthur, M. Cerezo, L. W. Harris, J. Hayhurst, C. Malangone, A. McMahon, J. Morales, E. Mountjoy, E. Sollis, D. Suveges, O. Vrousgou, P. L. Whetzel, R. Amode, J. A. Guillen, H. S. Riat, S. J. Trevanion, P. Hall, H. Junkins, P. Flicek, T. Burdett, L. A. Hindorff, F. Cunningham, H. Parkinson, The NHGRI-EBI GWAS Catalog of published genome-wide association studies, targeted arrays and summary statistics 2019. Nucleic Acids Res. 47, D1005–D1012 (2019).

107. M. Arnold, J. Raffler, A. Pfeufer, K. Suhre, G. Kastenmüller, SNiPA: an interactive, genetic variant-centered annotation browser. Bioinformatics. 31, 1334–1336 (2015).

108. X. Du, Y. Zhang, Q. Zhao, W. Qin, G. Ma, J. Fu, Q. Zhang, Effects of INSR genetic polymorphism on hippocampal volume and episodic memory in chinese type 2 diabetes. Acta Diabetol. 58, 1471–1480 (2021).

109. Y. Liu, S. Ran, Y. Lin, Y.-X. Zhang, X.-L. Yang, X.-T. Wei, Z.-X. Jiang, X. He, H. Zhang, G.-J. Feng, H. Shen, Q. Tian, H.-W. Deng, L. Zhang, Y.-F. Pei, Four pleiotropic loci associated with fat mass and lean mass. Int. J. Obes. . 44, 2113–2123 (2020).

110. L. O. Huang, A. Rauch, E. Mazzaferro, M. Preuss, S. Carobbio, C. S. Bayrak, N. Chami, Z. Wang, U. M. Schick, N. Yang, Y. Itan, A. Vidal-Puig, M. den Hoed, S. Mandrup, T. O. Kilpeläinen, R. J. F. Loos, Genome-wide discovery of genetic loci that uncouple excess adiposity from its comorbidities. Nat Metab. 3, 228–243 (2021).

111. S. Torkamandi, M. Bastami, H. Ghaedi, F. Moghadam, R. Mirfakhraie, M. D. Omrani, MAP3K1 May be a Promising Susceptibility Gene for Type 2 Diabetes Mellitus in an Iranian Population. Int J Mol Cell Med. 5, 134–140 (2016).

112. A. Al Shamsi, N. Al Hassani, M. Hamchou, R. Almazrouei, A. Mhanni, A novel missense heterozygous mutation in MAP3K1 gene causes 46, XY disorder of sex development: case report and literature review. Mol. Genet. Genomic Med. 8, e1514 (2020).

113. V. Todorovová, J. Hubacek, L. Dlouha, V. Adamkova, J. Pitha, M. Satny, R. Ceska, M. Vrablik, APOA5, GCKR, LRP1 AND MAP3K1 polymorphisms and the risk of acute coronary syndrome in the Czech population. Atherosclerosis. 331, e215 (2021).

114. A. Wesolowska-Andersen, G. Zhuo Yu, V. Nylander, F. Abaitua, M. Thurner, J. M. Torres, A. Mahajan, A. L. Gloyn, M. I. McCarthy, Deep learning models predict regulatory variants in pancreatic islets and refine type 2 diabetes association signals. Elife. 9 (2020), doi:10.7554/eLife.51503.

115. N. L. Harvey, R. S. Srinivasan, M. E. Dillard, N. C. Johnson, M. H. Witte, K. Boyd, M. W. Sleeman, G. Oliver, Lymphatic vascular defects promoted by Prox1 haploinsufficiency cause adult-onset obesity. Nat. Genet. 37, 1072–1081 (2005).

116. C. Schnoz, S. Moser, D. V. Kratschmar, A. Odermatt, D. Loffing-Cueni, J. Loffing, Deletion of the transcription factor Prox-1 specifically in the renal distal convoluted tubule causes hypomagnesemia via reduced expression of TRPM6 and NCC. Pflugers Arch. 473, 79–93 (2021).

117. L. M. M. Gommers, J. G. J. Hoenderop, R. J. M. Bindels, J. H. F. de Baaij, Hypomagnesemia in Type 2 Diabetes: A Vicious Circle? Diabetes. 65, 3–13 (2016).

118. L. J. Oost, J. I. P. van Heck, C. J. Tack, J. H. F. de Baaij, The association between hypomagnesemia and poor glycaemic control in type 1 diabetes is limited to insulin resistant individuals. Sci. Rep. 12, 1–7 (2022).

119. L. Kouřimská, A. Adámková, Nutritional and sensory quality of edible insects. NFS Journal. 4, 22–26 (2016).

120. M. Baron, A. Veres, S. L. Wolock, A. L. Faust, R. Gaujoux, A. Vetere, J. H. Ryu, B. K. Wagner, S. S. Shen-Orr, A. M. Klein, D. A. Melton, I. Yanai, A Single-Cell Transcriptomic Map of the Human and Mouse Pancreas Reveals Inter- and Intra-cell Population Structure. Cell Systems. 3, 346–360.e4 (2016).

121. L. Tosti, Y. Hang, O. Debnath, S. Tiesmeyer, T. Trefzer, K. Steiger, F. W. Ten, S. Lukassen, S. Ballke, A. A. Kühl, S. Spieckermann, R. Bottino, N. Ishaque, W. Weichert, S. K. Kim, R. Eils, C. Conrad, Single-Nucleus and In Situ RNA–Sequencing Reveal Cell Topographies in the Human Pancreas. Gastroenterology. 160, 1330–1344.e11 (2021).

122. M. Perez-Frances, L. van Gurp, M. V. Abate, V. Cigliola, K. Furuyama, E. Bru-Tari, D. Oropeza, T. Carreaux, Y. Fujitani, F. Thorel, P. L. Herrera, Pancreatic Ppy-expressing γ-cells display mixed phenotypic traits and the adaptive plasticity to engage insulin production. Nat. Commun. 12, 4458 (2021).

123. D. Grün, M. J. Muraro, J.-C. Boisset, K. Wiebrands, A. Lyubimova, G. Dharmadhikari, M. van den Born, J. van Es, E. Jansen, H. Clevers, E. J. P. de Koning, A. van Oudenaarden, De Novo Prediction of Stem Cell Identity using Single-Cell Transcriptome Data. Cell Stem Cell. 19, 266–277 (2016).

124. M. J. Muraro, G. Dharmadhikari, D. Grün, N. Groen, T. Dielen, E. Jansen, L. van Gurp, M. A. Engelse, F. Carlotti, E. J. P. de Koning, A. van Oudenaarden, A Single-Cell Transcriptome Atlas of the Human Pancreas. Cell Syst. 3, 385–394.e3 (2016).

125. Å. Segerstolpe, A. Palasantza, P. Eliasson, E.-M. Andersson, A.-C. Andréasson, X. Sun, S. Picelli, A. Sabirsh, M. Clausen, M. K. Bjursell, D. M. Smith, M. Kasper, C. Ämmälä, R. Sandberg, Single-Cell Transcriptome Profiling of Human Pancreatic Islets in Health and Type 2 Diabetes. Cell Metab. 24, 593–607 (2016).

126. N. Lawlor, J. George, M. Bolisetty, R. Kursawe, L. Sun, V. Sivakamasundari, I. Kycia, P. Robson, M. L. Stitzel, Single-cell transcriptomes identify human islet cell signatures and reveal cell-type-specific expression changes in type 2 diabetes. Genome Res. 27, 208–222 (2017).

127. Y. Xin, C. Adler, J. Kim, Y. Ding, M. Ni, Y. Wei, L. Macdonald, H. Okamoto, Single-cell RNA Sequencing and Analysis of Human Pancreatic Islets. J. Vis. Exp. (2019), doi:10.3791/59866.

128. M. M. Cooley, E. K. Jones, F. S. Gorelick, G. E. Groblewski, Pancreatic acinar cell protein synthesis, intracellular transport, and export. Pancreapedia: Exocrine Pancreas Knowledge Base (2020), doi:10.3998/panc.2020.15.

129. A. Grapin-Botton, Ductal cells of the pancreas. Int. J. Biochem. Cell Biol. 37, 504–510 (2005).

130. J. Maléth, P. Hegyi, Calcium signaling in pancreatic ductal epithelial cells: an old friend and a nasty enemy. Cell Calcium. 55, 337–345 (2014).

131. M. Priyadarshini, B. T. Layden, FFAR3 modulates insulin secretion and global gene expression in mouse islets. Islets. 7, e1045182 (2015).

132. C. Tang, K. Ahmed, A. Gille, S. Lu, H.-J. Gröne, S. Tunaru, S. Offermanns, Loss of FFA2 and FFA3 increases insulin secretion and improves glucose tolerance in type 2 diabetes. Nat. Med. 21, 173–177 (2015).

133. C. Berger, D. Zdzieblo, Glucose transporters in pancreatic islets. Pflugers Arch. 472, 1249–1272 (2020).

134. J. J. DiNicolantonio, J. H O’Keefe, Myo-inositol for insulin resistance, metabolic syndrome, polycystic ovary syndrome and gestational diabetes. Open Heart. 9 (2022), doi:10.1136/openhrt-2022-001989.

135. D. D. Moreno-Santillán, C. Machain-Williams, G. Hernández-Montes, J. Ortega, De Novo Transcriptome Assembly and Functional Annotation in Five Species of Bats. Sci. Rep. 9, 6222 (2019).

136. M. J. Coady, B. Wallendorff, D. G. Gagnon, J.-Y. Lapointe, Identification of a novel Na+/myo-inositol cotransporter. J. Biol. Chem. 277, 35219–35224 (2002).

137. M. A. Atkinson, M. Campbell-Thompson, I. Kusmartseva, K. H. Kaestner, Organisation of the human pancreas in health and in diabetes. Diabetologia. 63, 1966–1973 (2020).

138. Y. Xin, J. Kim, H. Okamoto, M. Ni, Y. Wei, C. Adler, A. J. Murphy, G. D. Yancopoulos, C. Lin, J. Gromada, RNA Sequencing of Single Human Islet Cells Reveals Type 2 Diabetes Genes. Cell Metab. 24, 608–615 (2016).

139. E. Bosi, L. Marselli, C. De Luca, M. Suleiman, M. Tesi, M. Ibberson, D. L. Eizirik, M. Cnop, P. Marchetti, Integration of single-cell datasets reveals novel transcriptomic signatures of β-cells in human type 2 diabetes. NAR Genom Bioinform. 2, lqaa097 (2020).

140. Y.-T. Chen, W.-D. Lin, W.-L. Liao, Y.-J. Lin, J.-G. Chang, F.-J. Tsai, PTPRD silencing by DNA hypermethylation decreases insulin receptor signaling and leads to type 2 diabetes. Oncotarget. 6, 12997–13005 (2015).

141. Y. Kang, H. Huang, H. Li, W. Sun, C. Zhang, Functional genetic variants in the 3’UTR of PTPRD associated with the risk of gestational diabetes mellitus. Exp. Ther. Med. 21, 562 (2021).

142. L. Stoll, J. Sobel, A. Rodriguez-Trejo, C. Guay, K. Lee, M. T. Venø, J. Kjems, D. R. Laybutt, R. Regazzi, Circular RNAs as novel regulators of β-cell functions in normal and disease conditions. Mol Metab. 9, 69–83 (2018).

143. S. Elumalai, U. Karunakaran, J.-S. Moon, K.-C. Won, NADPH Oxidase (NOX) Targeting in Diabetes: A Special Emphasis on Pancreatic β-Cell Dysfunction. Cells. 10 (2021), doi:10.3390/cells10071573.

144. T. Koufakis, A. Sertedaki, E.-B. Tatsi, C.-M. Trakatelli, S. N. Karras, E. Manthou, C. Kanaka-Gantenbein, K. Kotsa, First Report of Diabetes Phenotype due to a Loss-of-Function *ABCC8* Mutation Previously Known to Cause Congenital Hyperinsulinism. Case Rep. Genet. 2019, 3654618 (2019).

145. E. Araki, M. A. Lipes, M. E. Patti, J. C. Brüning, B. Haag 3rd, R. S. Johnson, C. R. Kahn, Alternative pathway of insulin signalling in mice with targeted disruption of the IRS-1 gene. Nature. 372, 186–190 (1994).

146. M. G. De Vas, J. L. Kopp, C. Heliot, M. Sander, S. Cereghini, C. Haumaitre, Hnf1b controls pancreas morphogenesis and the generation of Ngn3+ endocrine progenitors. Development. 142, 871–882 (2015).

147. C.-K. Kim, P. He, A. B. Bialkowska, V. W. Yang, SP and KLF Transcription Factors in Digestive Physiology and Diseases. Gastroenterology. 152, 1845–1875 (2017).

148. H. Kaneto, T.-A. Matsuoka, Role of pancreatic transcription factors in maintenance of mature β-cell function. Int. J. Mol. Sci. 16, 6281–6297 (2015).

149. H. J. Li, A. Kapoor, M. Giel-Moloney, G. Rindi, A. B. Leiter, Notch signaling differentially regulates the cell fate of early endocrine precursor cells and their maturing descendants in the mouse pancreas and intestine. Dev. Biol. 371, 156–169 (2012).

150. S. Spohrer, R. Groß, L. Nalbach, L. Schwind, H. Stumpf, M. D. Menger, E. Ampofo, M. Montenarh, C. Götz, Functional interplay between the transcription factors USF1 and PDX-1 and protein kinase CK2 in pancreatic β-cells. Sci. Rep. 7, 16367 (2017).

151. A. P. Sanchez, J. Zhao, Y. You, A.-E. Declèves, M. Diamond-Stanic, K. Sharma, Role of the USF1 transcription factor in diabetic kidney disease. Am. J. Physiol. Renal Physiol. 301, F271–9 (2011).

152. J. E. Gunton, R. N. Kulkarni, S. Yim, T. Okada, W. J. Hawthorne, Y.-H. Tseng, R. S. Roberson, C. Ricordi, P. J. O’Connell, F. J. Gonzalez, C. R. Kahn, Loss of ARNT/HIF1beta mediates altered gene expression and pancreatic-islet dysfunction in human type 2 diabetes. Cell. 122, 337–349 (2005).

153. H. S. Kang, K. Okamoto, Y.-S. Kim, Y. Takeda, C. D. Bortner, H. Dang, T. Wada, W. Xie, X.-P. Yang, G. Liao, A. M. Jetten, Nuclear orphan receptor TAK1/TR4-deficient mice are protected against obesity-linked inflammation, hepatic steatosis, and insulin resistance. Diabetes. 60, 177–188 (2011).

154. Y.-F. Lee, S. Liu, N.-C. Liu, R.-S. Wang, L.-M. Chen, W.-J. Lin, H.-J. Ting, H.-C. Ho, G. Li, E. J. Puzas, Q. Wu, C. Chang, Premature aging with impaired oxidative stress defense in mice lacking TR4. Am. J. Physiol. Endocrinol. Metab. 301, E91–8 (2011).

155. S. Wang, J. Skorczewski, X. Feng, L. Mei, J. E. Murphy-Ullrich, Glucose Up-regulates Thrombospondin 1 Gene Transcription and Transforming Growth Factor-β Activity through Antagonism of cGMP-dependent Protein Kinase Repression via Upstream Stimulatory Factor 2*. J. Biol. Chem. 279, 34311–34322 (2004).

156. L. Shi, S. Liu, D. Nikolic, S. Wang, High glucose levels upregulate upstream stimulatory factor 2 gene transcription in mesangial cells. J. Cell. Biochem. 103, 1952–1961 (2008).

157. R. K. Lex, W. Zhou, Z. Ji, K. N. Falkenstein, K. E. Schuler, K. E. Windsor, J. D. Kim, H. Ji, S. A. Vokes, GLI transcriptional repression is inert prior to Hedgehog pathway activation. Nat. Commun. 13, 808 (2022).

158. R. J. Weiss, P. N. Spahn, A. G. Toledo, A. W. T. Chiang, B. P. Kellman, J. Li, C. Benner, C. K. Glass, P. L. S. M. Gordts, N. E. Lewis, J. D. Esko, ZNF263 is a transcriptional regulator of heparin and heparan sulfate biosynthesis. Proceedings of the National Academy of Sciences. 117, 9311–9317 (2020).

159. C. Su, L. Gao, C. L. May, J. A. Pippin, K. Boehm, M. Lee, C. Liu, M. C. Pahl, M. L. Golson, A. Naji, the HPAP Consortium, S. F. A. Grant, A. D. Wells, K. H. Kaestner, The three-dimensional chromatin structure of the major human pancreatic cell types reveals lineage-specific regulatory architecture of T2D risk. bioRxiv (2022), p. 2021.11.30.470653.

160. M. Heddad Masson, C. Poisson, A. Guérardel, A. Mamin, J. Philippe, Y. Gosmain, Foxa1 and Foxa2 regulate α-cell differentiation, glucagon biosynthesis, and secretion. Endocrinology. 155, 3781–3792 (2014).

161. S. L. Park, I. Cheng, S. A. Pendergrass, A. M. Kucharska-Newton, U. Lim, J. L. Ambite, C. P. Caberto, K. R. Monroe, F. Schumacher, L. A. Hindorff, M. T. Oetjens, S. Wilson, R. J. Goodloe, S.-A. Love, B. E. Henderson, L. N. Kolonel, C. A. Haiman, D. C. Crawford, K. E. North, G. Heiss, M. D. Ritchie, L. R. Wilkens, L. Le Marchand, Association of the FTO obesity risk variant rs8050136 with percentage of energy intake from fat in multiple racial/ethnic populations: the PAGE study. Am. J. Epidemiol. 178, 780–790 (2013).

162. T. Bego, A. Čaušević, T. Dujić, M. Malenica, Z. Velija-Asimi, B. Prnjavorac, J. Marc, J. Nekvindová, V. Palička, S. Semiz, Association of FTO Gene Variant (rs8050136) with Type 2 Diabetes and Markers of Obesity, Glycaemic Control and Inflammation. J. Med. Biochem. 38, 153–163 (2019).

163. H. Jia, L. Yu, Z. Jiang, Q. Ji, Association between IGF2BP2 rs4402960 polymorphism and risk of type 2 diabetes mellitus: a meta-analysis. Arch. Med. Res. 42, 361–367 (2011).

164. D. El-Lebedy, I. Ashmawy, A. A. Ibrahim, Common Variants in IGF2BP2 Gene rs4402960 and rs1470579 Polymorphisms Associate with Type 2 Diabetes Mellitus in Egyptians: A Replication Study. International Journal of Diabetes Research. 4, 43–48 (2015).

165. M.-Q. Mo, L. Pan, Q.-M. Lu, Q.-L. Li, Y.-H. Liao, The association of the CMIP rs16955379 polymorphism with dyslipidemia and the clinicopathological features of IgA nephropathy. Int. J. Clin. Exp. Pathol. 11, 5008–5023 (2018).

166. L. Pan, Y.-H. Liao, M.-Q. Mo, Q.-H. Zhang, R.-X. Yin, CMIP SNPs and their haplotypes are associated with dyslipidaemia and clinicopathologic features of IgA nephropathy. Biosci. Rep. 40 (2020), doi:10.1042/BSR20202628.

167. Z. B. Mehta, N. Fine, T. J. Pullen, M. C. Cane, M. Hu, P. Chabosseau, G. Meur, A. Velayos-Baeza, A. P. Monaco, L. Marselli, P. Marchetti, G. A. Rutter, Changes in the expression of the type 2 diabetes-associated gene VPS13C in the β-cell are associated with glucose intolerance in humans and mice. Am. J. Physiol. Endocrinol. Metab. 311, E488–507 (2016).

168. T. Kuo, M. J. Kraakman, M. Damle, R. Gill, M. A. Lazar, D. Accili, Identification of C2CD4A as a human diabetes susceptibility gene with a role in β cell insulin secretion. Proc. Natl. Acad. Sci. U. S. A. 116, 20033–20042 (2019).

169. R. D. Albanus, X. Tang, H. J. Taylor, N. Manickam, M. Erdos, N. Narisu, Y. Han, P. Orchard, A. Varshney, C. Liu, A. Naji, HPAP Consortium, F. S. Collins, S. Chen, S. C. J. Parker, Single-cell gene expression and chromatin accessibility profiling of human pancreatic islets at basal and stimulatory conditions nominates mechanisms of type 1 diabetes genetic risk. bioRxiv (2022), p. 2022.11.12.516291.

170. F. Wei, C. Cai, S. Feng, J. Lv, S. Li, B. Chang, H. Zhang, W. Shi, H. Han, C. Ling, P. Yu, Y. Chen, N. Sun, J. Tian, H. Jiao, F. Yang, M. Li, Y. Wang, L. Zou, L. Su, J. Li, R. Li, H. Qiu, J. Shi, S. Liu, M. Chang, J. Lin, L. Chen, W.-D. Li, TOX and CDKN2A/B Gene Polymorphisms Are Associated with Type 2 Diabetes in Han Chinese. Sci. Rep. 5, 11900 (2015).

171. G. Casotti, K. C. Richardson, J. S. Bradley, Ecomorphological constraints imposed by the kidney component measurements in honeyeater birds inhabiting different environments. J. Zool. . 231, 611–625 (1993).

172. G. Casotti, K. C. Richardson, A stereological analysis of kidney structure of honeyeater birds (Meliphagidae) inhabiting either arid or wet environments. J. Anat. 180 (Pt 2), 281– 288 (1992).

173. C. A. Beuchat, M. R. Preest, E. J. Braun, Glomerular and medullary architecture in the kidney of Anna’s Hummingbird. J. Morphol. 240, 95–100 (1999).

174. I. Bravo-Ruiz, M. Á. Medina, B. Martínez-Poveda, From Food to Genes: Transcriptional Regulation of Metabolism by Lipids and Carbohydrates. Nutrients. 13 (2021), doi:10.3390/nu13051513.

175. N. Matharu, N. Ahituv, Modulating gene regulation to treat genetic disorders. Nat. Rev. Drug Discov. 19, 757–775 (2020).

176. M. Bernt, A. Donath, F. Jühling, F. Externbrink, C. Florentz, G. Fritzsch, J. Pütz, M. Middendorf, P. F. Stadler, MITOS: improved de novo metazoan mitochondrial genome annotation. Mol. Phylogenet. Evol. 69, 313–319 (2013).

177. Y. Luo, B. C. Hitz, I. Gabdank, J. A. Hilton, M. S. Kagda, B. Lam, Z. Myers, P. Sud, J. Jou, K. Lin, U. K. Baymuradov, K. Graham, C. Litton, S. R. Miyasato, J. S. Strattan, O. Jolanki, J.-W. Lee, F. Y. Tanaka, P. Adenekan, E. O’Neill, J. M. Cherry, New developments on the Encyclopedia of DNA Elements (ENCODE) data portal. Nucleic Acids Res. 48, D882–D889 (2020).

178. C. Hafemeister, R. Satija, Normalization and variance stabilization of single-cell RNA-seq data using regularized negative binomial regression. Genome Biol. 20, 296 (2019).

179. H. Wickham, ggplot2: Elegant Graphics for Data Analysis (2016), (available at https://ggplot2.tidyverse.org).

180. D. Meyer, A. Zeileis, K. Hornik, The Strucplot Framework: Visualizing Multi-way Contingency Tables with vcd. J. Stat. Softw. 17, 1–48 (2007).

181. R. Kolde, pheatmap: Pretty heatmaps (Github; https://github.com/raivokolde/pheatmap).

182. C. A. Schneider, W. S. Rasband, K. W. Eliceiri, NIH Image to ImageJ: 25 years of image analysis. Nat. Methods. 9, 671–675 (2012).

183. D. R. Stirling, M. J. Swain-Bowden, A. M. Lucas, A. E. Carpenter, B. A. Cimini, A. Goodman, CellProfiler 4: improvements in speed, utility and usability. BMC Bioinformatics. 22, 433 (2021).

184. M. V. Kuleshov, M. R. Jones, A. D. Rouillard, N. F. Fernandez, Q. Duan, Z. Wang, S. Koplev, S. L. Jenkins, K. M. Jagodnik, A. Lachmann, M. G. McDermott, C. D. Monteiro, G. W. Gundersen, A. Ma’ayan, Enrichr: a comprehensive gene set enrichment analysis web server 2016 update. Nucleic Acids Res. 44, W90–7 (2016).

185. A. S. Hinrichs, D. Karolchik, R. Baertsch, G. P. Barber, G. Bejerano, H. Clawson, M. Diekhans, T. S. Furey, R. A. Harte, F. Hsu, J. Hillman-Jackson, R. M. Kuhn, J. S. Pedersen, A. Pohl, B. J. Raney, K. R. Rosenbloom, A. Siepel, K. E. Smith, C. W. Sugnet, A. Sultan-Qurraie, D. J. Thomas, H. Trumbower, R. J. Weber, M. Weirauch, A. S. Zweig, D. Haussler, W. J. Kent, The UCSC Genome Browser Database: update 2006. Nucleic Acids Res. 34, D590–8 (2006).

186. R. S. Harris, thesis, The Pennsylvania State University (2007).

187. T. L. Bailey, J. Johnson, C. E. Grant, W. S. Noble, The MEME Suite. Nucleic Acids Res. 43, W39–49 (2015).

188. J. A. Castro-Mondragon, R. Riudavets-Puig, I. Rauluseviciute, R. B. Lemma, L. Turchi, R. Blanc-Mathieu, J. Lucas, P. Boddie, A. Khan, N. Manosalva Pérez, O. Fornes, T. Y. Leung, A. Aguirre, F. Hammal, D. Schmelter, D. Baranasic, B. Ballester, A. Sandelin, B. Lenhard, K. Vandepoele, W. W. Wasserman, F. Parcy, A. Mathelier, JASPAR 2022: the 9th release of the open-access database of transcription factor binding profiles. Nucleic Acids Res. 50, D165–D173 (2022).

189. S. T. Sherry, M. H. Ward, M. Kholodov, J. Baker, L. Phan, E. M. Smigielski, K. Sirotkin, dbSNP: the NCBI database of genetic variation. Nucleic Acids Res. 29, 308–311 (2001).

